# Specialized late cingulo-opercular network activation elucidates the mechanisms underlying decisions about ambiguity

**DOI:** 10.1101/2023.02.25.529948

**Authors:** Jordan E. Pierce, Nathan M. Petro, Elizabeth Clancy, Caterina Gratton, Steven E. Petersen, Maital Neta

**Author notes:** corresponding author, address: B76 East Stadium, Lincoln, NE 68588.

## Abstract

Cortical task control networks, including the cingulo-opercular (CO) network play a key role in decision-making across a variety of functional domains. In particular, the CO network functions in a performance reporting capacity that supports successful task performance, especially in response to errors and ambiguity. In two studies testing the contribution of the CO network to ambiguity processing, we presented a valence bias task in which masked clearly and ambiguously valenced emotional expressions were slowly revealed over several seconds. This slow reveal task design provides a window into the decision-making mechanisms as they unfold over the course of a trial. In the main study, the slow reveal task was administered to 32 young adults in the fMRI environment and BOLD time courses were extracted from regions of interest in three control networks. In a follow-up study, the task was administered to a larger, online sample (n = 81) using a more extended slow reveal design with additional unmasking frames. Positive judgments of surprised faces were uniquely accompanied by slower response times and strong, late activation in the CO network. These results support the initial negativity hypothesis, which posits that the default response to ambiguity is negative and positive judgments are associated with a more effortful controlled process, and additionally suggests that this controlled process is mediated by the CO network. Moreover, ambiguous trials were characterized by a second CO response at the end of the trial, firmly placing CO function late in the decision-making process.

## 1. Introduction

Decision-making is a central cognitive process that is necessary for myriad daily functions that help us to navigate an ever-changing, complex world. Ambiguity – a pervasive characteristic of this complex world – arises when insufficient information is available for guiding our decisions and poses challenges for determining an optimal path forward. One line of neuroimaging research has explored the neural mechanisms of decision-making in the context of ambiguity using an extended perceptual task (Gratton et al., 2017; Neta et al., 2017). In this “slow reveal” paradigm (see also Ploran et al., 2007), individuals must identify an object that is unmasked slowly over several seconds or report a word that is revealed one letter at a time. An analysis of several such functional magnetic resonance imaging (fMRI) studies indicated that the cingulo-opercular (CO) and fronto-parietal (FP) control networks of the brain were recruited during decision-making and exhibited distinct temporal characteristics. Responses in these task control networks were differentiated by the onset (early vs. late) and duration (extended vs. transient) of the hemodynamic response (Gratton et al., 2017, see also Coste & Kleinschmidt, 2016; Dosenbach et al., 2008). Specifically, early activation in the left FP network supported evidence accumulation and increased throughout the trial, while later extended activation in the right FP network reflected processing after the response occurred. Finally, late, transient activation in the CO network corresponded to performance reporting at the end of a trial (Gratton et al., 2017).

The performance reporting function of the CO network – which includes regions such as the dorsal anterior cingulate cortex, medial prefrontal cortex, anterior insula, and frontal operculum – was investigated further using the same slow reveal paradigm for object identification with the addition of an ambiguity condition (Neta et al., 2017). Performance accuracy and brain activation were measured on trials showing objects with unique, easily identifiable outlines (e.g., butterfly) or objects with ambiguous, easily mistaken outlines (e.g., toothbrush) that were gradually unmasked. Ambiguous stimuli and error trials (self-reported misidentifications) yielded interactive effects in the late response of the CO network, suggesting overlapping but distinct information processing for these functions as related to performance reporting that is heightened in the context of these trial types (see also Neta et al., 2014). This is consistent with work demonstrating that the CO shows increased activation in response to ambiguity in emotional valence (Neta et al., 2013) and many other domains (Neta et al., 2014; Sterzer et al., 2002; Thompson-Schill et al., 1997). Based on these previous findings regarding the function of the CO network in processing ambiguity, this network was of particular interest in the current work on ambiguously valenced emotional expressions.

Emotional expressions afford the opportunity to explore responses to ambiguity using a familiar and salient stimulus that is frequently used to help us navigate our environment. Certain facial expressions express positive or negative emotional valence relatively clearly, such as a happy or fearful expression, and quickly convey important social cues about potential rewards or threats in the environment. Other expressions, however, may convey a signal that is ambiguous in valence, such as a surprised expression, which could be interpreted as either positive or negative depending on the context (Kim et al., 2004; Neta et al., 2011; Neta & Kim, 2022). Thus, if individuals are asked to evaluate surprised expressions without context, valence judgments reflect this ambiguity and range from consistently positive appraisals to a mix of positive and negative to consistently negative appraisals in a trait-like manner that is termed “valence bias” (Harp et al., 2022; Kim et al., 2003; Neta et al., 2021).

Much of the work on this topic over the last 15 years has come to focus on determining the underlying processes that support valence bias. This work has demonstrated that the initial response to ambiguity tends to be negative, and that positive judgments may require a slower, more effortful process that helps to overcome the initial negativity (i.e., the initial negativity hypothesis; Neta et al., 2021; Neta & Tong, 2016; Neta & Whalen, 2010). For example, the presentation of low spatial frequency images of facial expressions (that can be processed quickly by the visual system) biased responses to ambiguous trials towards negative judgments (Neta & Whalen, 2010). Another study demonstrated that when participants were encouraged to delay their response to ambiguity, their judgments shifted towards positivity (Neta & Tong, 2016). Finally, by using a mouse-tracking approach, studies revealed that only positive judgments of ambiguous stimuli are characterized by an initial motoric attraction to the competing (negative) response option (Brown et al., 2017; Neta et al., 2021). Notably, the additional processing time and response competition for positive judgments is thought to be related to a regulatory process that helps to overcome the initial negativity (Neta, 2024; Neta et al., 2022; Petro et al., 2018).

In the current study, the slow reveal paradigm was combined with the valence bias task to probe how cortical task control networks support the underlying decision-making process during emotional valence judgments of ambiguity and, specifically, to test the initial negativity hypothesis that positive judgments would necessitate greater control. Participants were shown facial expressions with non-ambiguous valence – happy, angry, and fearful expressions – and ambiguous valence – surprised expressions. The faces were slowly unmasked and participants were instructed to categorize each face as expressing positive or negative valence. Perceptual uncertainty may impact recognition of certain facial expressions as a function of the similarity of morphological features, potentially resulting in slower or inaccurate judgments of their emotional valence. Importantly, the valence ambiguity inherent in a surprised expression was predicted to contribute to further uncertainty – slowing responses for this condition. These slower responses then might allow participants the opportunity to engage in a regulatory process that not only yields more positive surprised valence ratings, but also helps to identify the brain networks that support this regulatory process. Given the prevalent decision-making role for CO in response to ambiguity across many different domains of scientific inquiry (Neta et al., 2013, 2014; Poudel et al., 2020; Sterzer et al., 2002; Thompson-Schill et al., 1997), we hypothesized that the CO network would show greater activation during judgments of ambiguously valenced facial expressions and, more specifically, show greater and more extended activation when participants make more positive judgments of ambiguity.

Finally, we attempted to replicate and extend the behavioral findings from the MRI study in a follow-up online study that used a more extended unmasking phase. Specifically, an increased number of frames (i.e., smaller steps between frames) during the slow reveal was intended to provide greater sensitivity to differences in the relative timing of the valence decision-making process for ambiguous trials. Given that positive judgments of surprised faces are often slower and putatively require overriding the default negativity, we predicted that when positive judgments occurred, they would be made later in the unmasking process than negative judgments.

## 2. Methods

### 2.1 Participants

For the MRI study, thirty-seven participants were recruited from the Lincoln community via publicly posted flyers. Participants reported no history of neurological or psychiatric disorders and no use of psychotropic medications, were right-handed, had normal or corrected-to-normal vision, and were naive to the purpose of the study. One participant could not complete the MRI scan due to an unremovable piercing and four participants data were excluded due to technical issues. The final sample included 32 participants (20 female/12 male, mean age = 20.6 years (SD=1.7)). The target sample size was determined based on sample sizes in other slow reveal fMRI studies leveraging within-subject analyses (N=27, Neta et al., 2017; N=13, Ploran et al., 2007) and a power analysis using effect sizes from data reported in Neta et al., 2017, which used a slow reveal paradigm to examine cingulo-opercular responses to perceptual ambiguity. Our analyses focus on contrasts between ambiguous and clear valence expressions, and so we based our power analyses on a repeated measures ANOVA to ensure that we have sufficient power to discriminate the within-subject emotion conditions. The power analysis was conducted in G*Power (Faul et al., 2007) and indicated that with an *η*^2^ = .21, a repeated measures ANOVA requires 8 participants, but a bivariate correlation (with behavior) with r = 0.5 requires 29 participants to achieve 80% power to detect effects. All participants provided written informed consent and all procedures were approved by the Institutional Review Board at the University of Nebraska-Lincoln.

### 2.2 Procedure

Participants received verbal task instructions prior to entering the MRI scanner, then were positioned on their back with padding to secure their head in a comfortable, static position where they could view the experimental paradigm display via a mirror attached to the head coil. Responses were collected via an MRI-compatible button box with the right index and middle finger (with counterbalanced assignment to response choices). At the start of the functional scans, task instructions were presented on the screen and participants were given a verbal reminder. Participants were instructed to categorize each face as either positive or negative twice during the trial: once as the picture was being unmasked (“noise” condition) – as soon as they had a reasonably confident judgment – and again when the picture was fully revealed (“reveal” condition). They were told that the two responses could be the same or different. All participants then completed three practice trials containing angry and happy faces as examples of negative and positive valence (no ambiguous examples were given), with identities that were not used in the main experiment. Following the practice trials, the participants were given the opportunity to ask for clarification before beginning the task.

### 2.3 Task Design

On each trial, a grayscale face was occluded by a black mask that was slowly degraded over the course of five 2-second frames (**Figure 1**). Each trial began with a 2 sec black screen, then the black mask was degraded by randomly removing pixels following a Gaussian distribution over the five “noise” frames such that 0.75, 1.25, 3.0, 7.0, and 14.0% of the face image was visible at each successive step. Each trial ended with the presentation of the fully revealed face stimulus for 2 seconds. Thus, the duration of each trial was 14 seconds. During the inter-trial interval (ITI), a fixation dot was presented at the center of the screen for 2, 4, or 6 seconds (equally presented across trial conditions).

**Figure 1.**
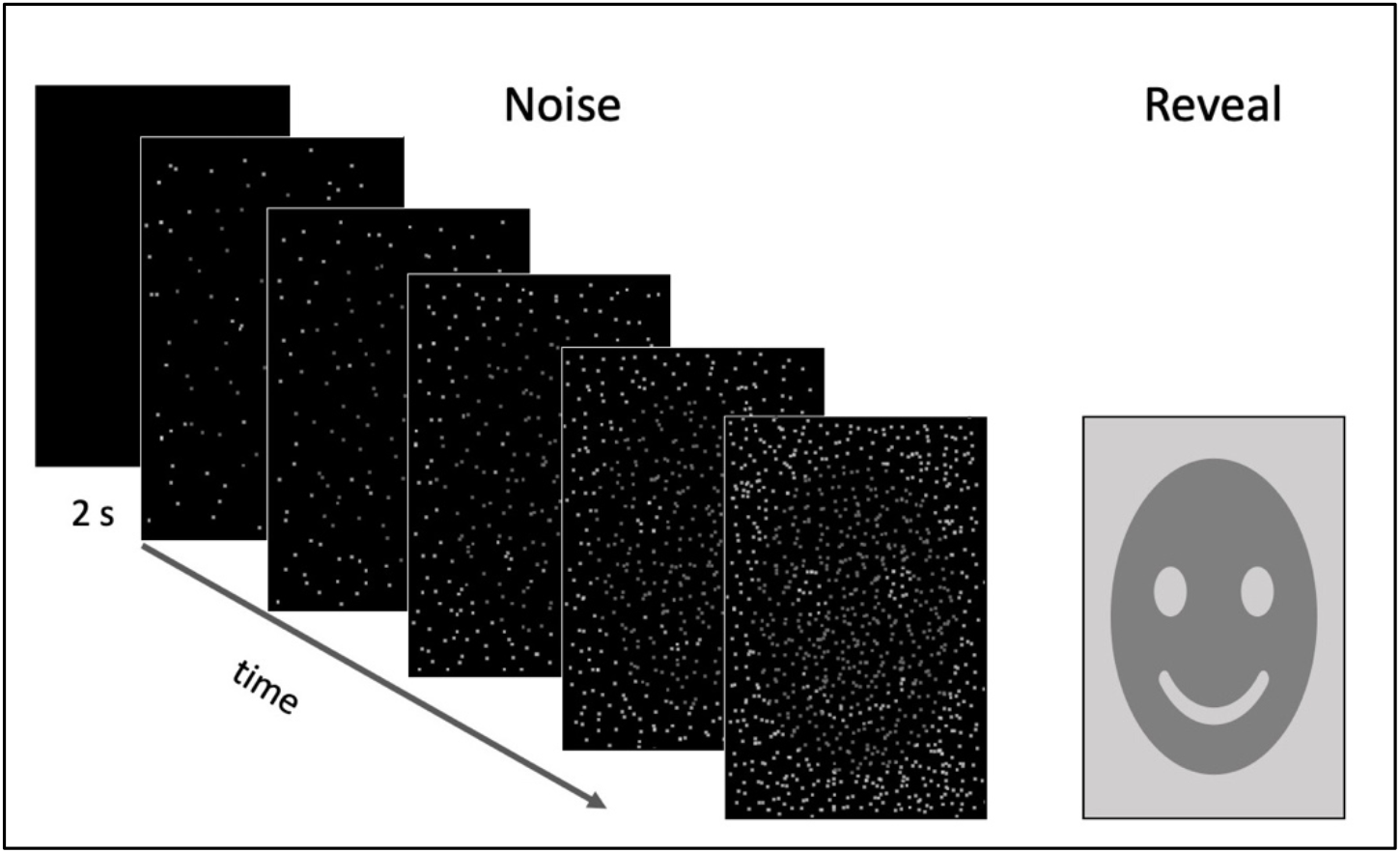
Task design for the slow reveal of emotional facial expressions. In the main study, each trial lasted 14 seconds, consisting of seven 2-second frames: one black frame, five “noise” frames with a decreasing number of masking pixels, and a final “reveal” frame. In between trials, a fixation dot appeared for a variable ITI of 2, 4, or 6 seconds. Here a smiley icon is used in place of the photograph shown to participants.

There was a total of 240 trials, with an equal distribution of faces displaying a happy, angry, fearful, or surprised expression (60 unique examples of each) across ten separate runs. Given that temporal context can shift responses to surprised faces (Neta et al., 2011), expressions were presented in pseudorandom order such that the surprised faces were preceded by an equal number of clearly positive and negative faces. The face stimuli were drawn from the NimStim Set of Facial Expressions (Tottenham et al., 2009), the Karolinska Directed Emotional Faces database (Goeleven et al., 2008), the Umea University Database of Facial Expressions, (Samuelsson et al., 2012), and the Ekman Pictures of Facial Affect (Ekman, 1976). Each model contributed one to four emotional expression images and no image was presented more than once during the task. The luminance of all faces was equated using the SHINE toolbox (Willenbockel et al., 2010) for MATLAB (Mathworks Inc, Natick, MA).

### 2.4 MRI Parameters

The MRI data were collected at the University of Nebraska-Lincoln Center for Brain, Biology, & Behavior on a Siemens 3T Skyra scanner using a 32-channel head coil. Structural images were acquired using a T1-weighted 3D multi-echo MPRAGE sequence (TR = 2530 ms, TE = 1.69 ms, 4 averages, slices = 176 interleaved, voxel size = 1.0 mm^3^, matrix = 256 × 256, FOV = 256 mm, flip angle = 7°, total acquisition time = 6:00). Blood oxygen level-dependent (BOLD) activity was measured using a multi-band EPI scanning sequence (TR = 1000 ms, TE = 29.8 ms, multi-band factor = 3, slices = 51 interleaved, voxel size = 2.50 mm^3^, matrix = 84 × 84 mm, FOV = 168 mm, flip angle = 60°, 454 volumes, total acquisition time = 7:48 per run), and slices were acquired parallel to the AC-PC plane, positioned to cover the entire brain.

### 2.5 FMRI Study Behavioral Analysis

Average valence judgments for each emotion were calculated as the number of negative responses out of the number of total trials for each participant, multiplied by 100 (i.e., percent negative). Average judgments were entered into a linear mixed model with fixed effects of emotion (4 levels – happy, angry, fearful, surprised), trial phase (2 levels – noise, reveal), and their interaction, with a random intercept by subject. Emotion and phase were modeled as repeated measures with a diagonal covariance structure and Satterthwaite estimates were used for determining degrees of freedom for the fixed effects. Subsequently, incorrect judgments for clearly valenced emotional expressions (happy judged as negative and angry or fearful judged as positive) were considered errors and not included in the response time analyses; for surprised faces, both positive and negative judgments were retained as separate conditions.

The average response frame (out of 5) was calculated for each of five emotion conditions (happy, angry, fearful, positive surprised, and negative surprised) for each participant. Response time (RT) was calculated for responses in both the noise phase and the reveal phase. The noise RT was calculated from the beginning of the first noise frame, giving a maximum possible value of 10 seconds (2 sec x 5 frames), while the reveal RT had a maximum possible value of 2 seconds. Noise and reveal RTs were entered into separate linear mixed models with fixed effects of emotion (5 levels) and a random intercept by subject. Emotion was modeled as a repeated measure with a diagonal covariance structure and Satterthwaite estimates were used for determining degrees of freedom for the fixed effects. All statistical analyses were conducted in SPSS version 28 (IBM, Cary, NC, USA).

## 2.6 FMRI Data Analysis

### 2.6.1 Preprocessing

Functional data were analyzed using the AFNI software package (Cox, 1996, 2012). Preprocessing consisted of de-spiking of time series outliers, slice timing correction, alignment of functional volumes to each other and the individual anatomical image, standardization to the Talairach atlas space (Talairach & Tournoux, 1988), smoothing with a 6-mm FWHM kernel, and scaling of each voxel to a mean of 100 to convert arbitrary units to percent signal change. Volumes with a motion shift where the Euclidean norm of the derivative was greater than 0.3 were censored (0.45% of volumes removed across all participants).

### 2.6.2 HRF Time Courses by Emotional Expression

Preprocessed functional data were entered into a general linear model in which individual responses were sorted by the emotion shown and valence judgment made (see Supplemental Material and Figure S1 for ROI analysis combined across all emotions). Thus, there were regressors for positive judgments of happy faces, negative judgments of angry faces, negative judgments of fearful faces, positive judgments of surprised faces, and negative judgments of surprised faces, combined across response frames 2-5. The “TENT” function was used to estimate the amplitude of the hemodynamic response at each TR from 0-21 seconds after stimulus onset without assuming a predetermined shape for the HRF in each voxel. Regressors of no interest included error trials (negative judgments of happy faces and positive judgments of angry and fearful faces), anticipatory responses (frame 1 – when there was insufficient visual information to make a decision), polynomials for each run (two terms), and six motion parameters estimated during alignment (x, y, z shift/rotation).

Subsequently, the beta values at each time point (i.e., the estimated hemodynamic response function (HRF)) were extracted from 16 regions of interest (ROIs; **Figure 2**) based on a previous study using a slow reveal task (Gratton et al., 2017). Each ROI was defined as a 5-mm sphere centered on the peak coordinates of each region. Following the approach of Gratton et al., 2017, data from these 16 ROIs were combined into three ROI clusters: cingulo-opercular (CO), left frontoparietal (L FP), and right frontoparietal (R FP). The CO cluster consisted of four ROIs (as labelled in Gratton et al., 2017): left anterior insula/frontal operculum (AI/FO), right AI/FO, pre-supplemental motor area (SMA) 1, and pre-SMA 2. The L FP cluster consisted of eight ROIs: left inferior parietal sulcus (IPS) 1, left IPS 2, left IPS 3, left frontal, left middle frontal gyrus (MFG) 1, left MFG 2, right IPS 2, and right frontal. The R FP cluster consisted of four ROIs: right IPS 1, right inferior frontal gyrus (IFG), right MFG 1, and right MFG 2. Additionally, responses from 5 visual ROIs based on another slow reveal study (Ploran et al., 2007) were averaged together in a “Sensory” ROI cluster as a control condition in which no differences were expected by emotion (see Supplemental Table S1 for all regions and their coordinates).

**Figure 2.**
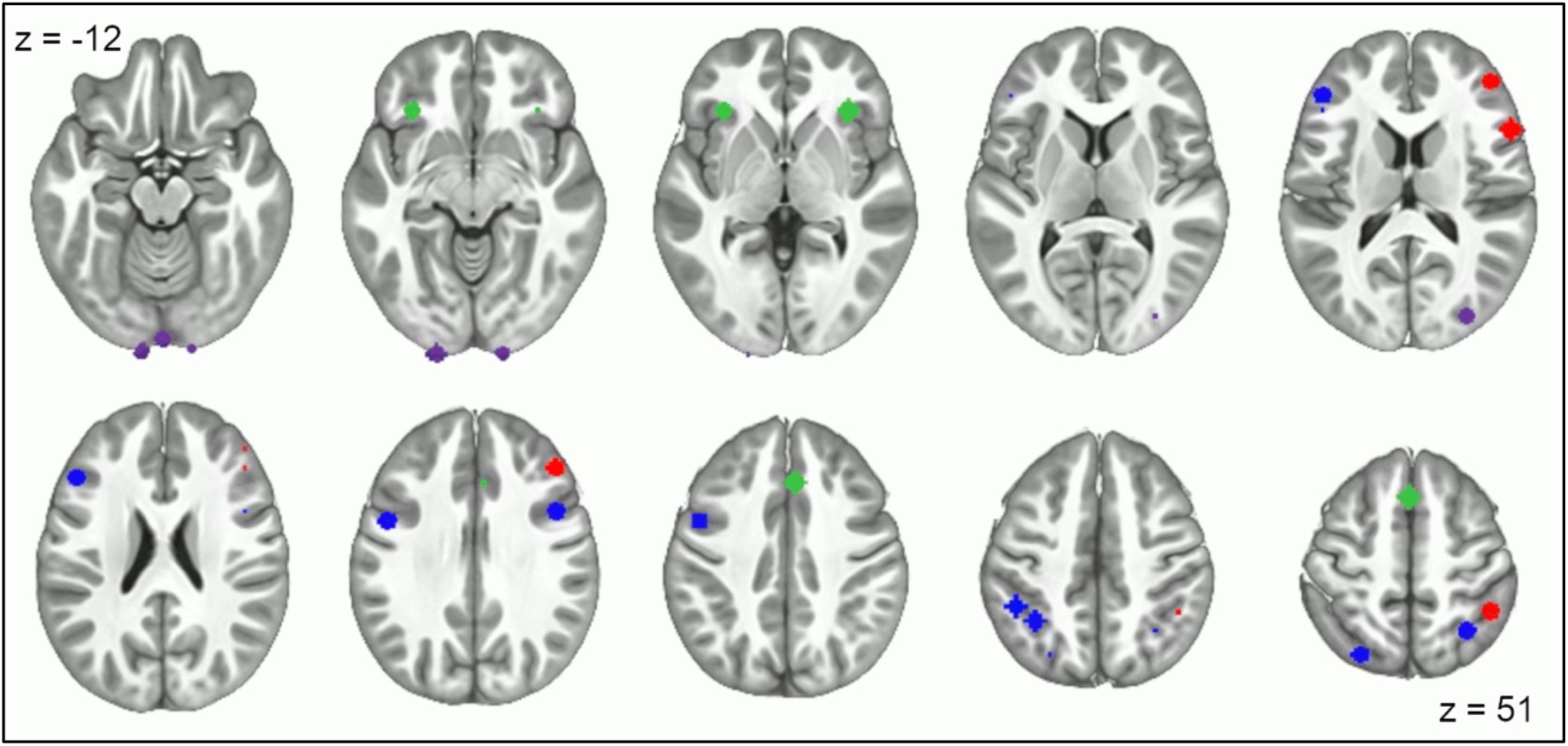
Location of the ROIs extracted based on Gratton et al., 2017 and Ploran et al, 2007. Colors indicate the ROI cluster: CO (green), left FP (blue), right FP (red), and Sensory (purple). Coordinates for each ROI are provided in Supplemental Table S1.

To compare the responses for each emotion, beta values were entered into a linear mixed model with fixed effects of emotion (5 levels), time (11 timepoints; with linear and quadratic terms), and their interaction (emotion X linear time), with a random intercept by subject. For this analysis, only the second half of the HRF time course was examined (11-21 seconds) in order to focus on the period that contains the hemodynamic peak (see Figure 3) and exclude early time points with minimal activation across conditions. Given the behavioral differences between emotional expressions, RT from the reveal phase was also included as a covariate. Emotion and time were modeled as repeated measures with a diagonal covariance structure and Satterthwaite estimates were used for determining degrees of freedom for the fixed effects. Separate models were fit for each of the three ROI clusters of interest (CO, L FP, and R FP) and the control cluster (Sensory). For positive surprised trials, 10 participants were removed due to a low trial count (< 4 surprised trials with a “positive” response, see Supplemental Table S2; remaining N=22); all other emotion conditions included all participants (N=32).

**Figure 3.**
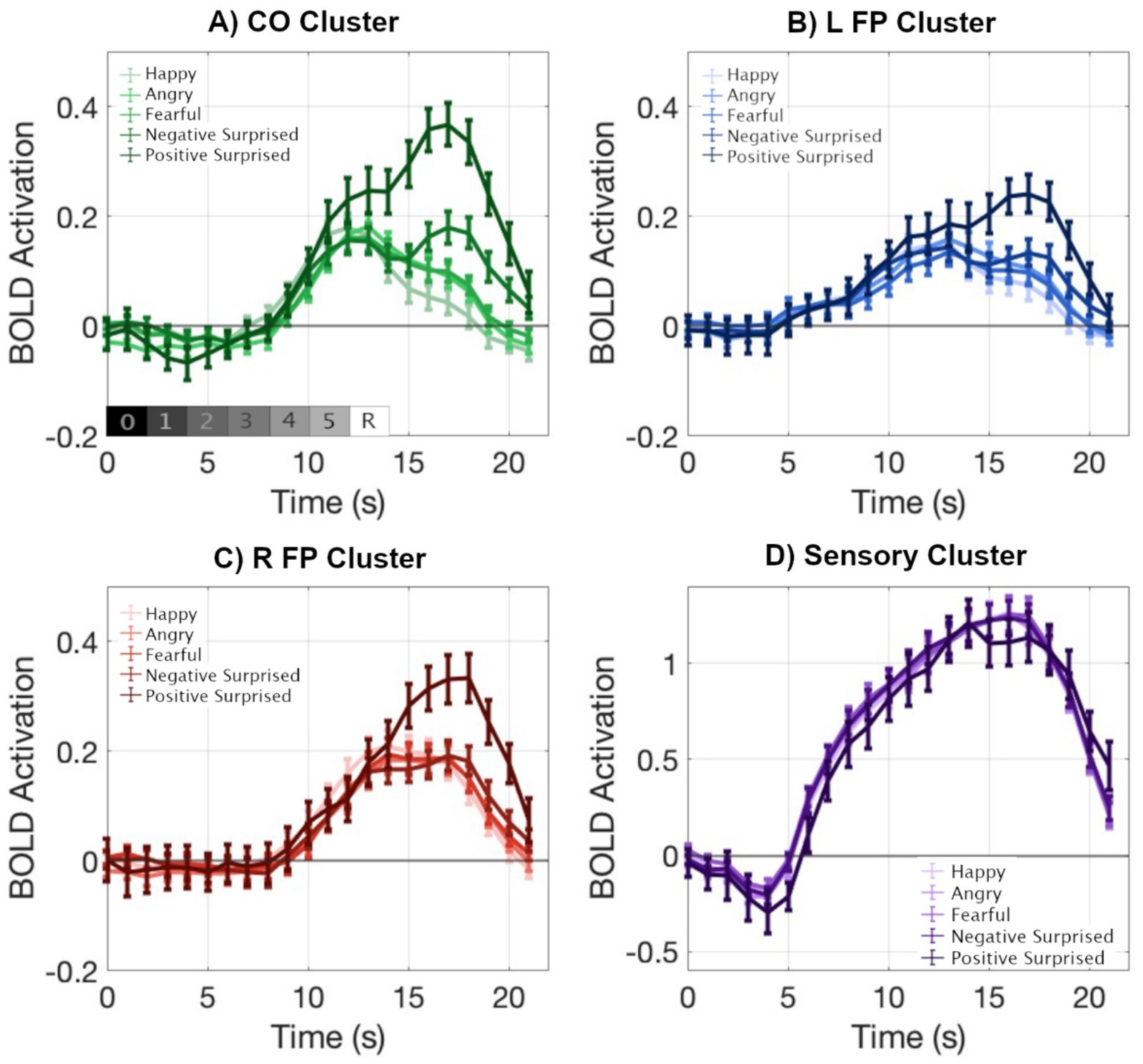
Estimated HRF time courses by emotional expression in each of the ROI clusters. Emotions included correct clear valence trials (happy judged as positive, angry/fearful judged as negative) and ambiguous valence (surprised) trials judged as either negative or positive. In all three clusters of interest, positive surprised trials yielded the strongest response in the last half of the HRF; there were no effects of emotion in the control (Sensory) cluster. The gray bar represents the timing of each frame of the trial from 0 = black screen to 5 = final noise frame and R = fully revealed image.

### 2.6.3 Late CO Activation and Behavior

From the initial analysis, there was a second, late peak observed in the CO responses to surprised faces (see Fig. 3) that we predicted might reflect a response to the final reveal of the (still ambiguously valenced) face. To explore this possibility, beta values were extracted for each emotional expression for the period from 16-19 seconds in the trial, corresponding to the period 4-7 seconds after the final reveal image was displayed (i.e., when an HRF peak for the reveal image would occur). The average activation in this period was correlated with participants’ average reveal RT for each emotion to determine if CO activation was dependent on RT differences. Additionally, the activation for positive and negative surprised trials was correlated with participants’ overall valence bias (percent of negative judgments) for the reveal phase to determine if CO activation was driven by extended processing for positive judgments.

### 2.6.4 HRF Time Courses as a Function of Switching versus Staying

To further examine the differences in HRF time courses by emotional expression, we conducted an exploratory analysis to compare trials on which participants made the same behavioral judgment during the noise and reveal phases of the trial and trials on which participants switched their response during the reveal phase. The latter “switch” condition included clear valence trials on which participants made an error, presumably recognized the error, and corrected their response when seeing the fully revealed face. Importantly, the “switch” condition also included ambiguous trials (which do not have an objectively accurate response) on which participants changed their response at the reveal phase, perhaps because they decided that their initial judgment was inappropriate. Conversely, the “stay” condition included only trials on which participants did not change their valence judgment, presumably because they were sufficiently confident in their initial response and did not consider it an error.

For this analysis, clear valence errors (e.g., negative judgment of a happy face) were excluded for the stay condition and clear valence correct trials (e.g., positive judgment of a happy face) were excluded for the switch condition given an insufficient number of trials (< two trials on average). Additionally, within each of the remaining conditions, individual participants were excluded if they had fewer than three trials in a particular response condition (see Supplemental Table S3 for trial counts by condition). This requirement led to a different number of participants being included across conditions (<u>stay</u> condition: positive happy/negative angry/negative fearful/negative surprised: N=32; positive surprised: N=17; <u>switch</u> condition: negative happy: N=15; positive angry: N=24; positive fearful: N=31; positive surprised: N=14; negative surprised: N=19). Nonetheless, this approach provides the best estimate of the HRFs using the maximum amount of available data for this initial exploratory analysis of potential factors contributing to the HRF differences by emotion.

As with the prior analysis by emotional expression, a linear mixed model was fit with fixed effects of emotion (5 levels), time (11 timepoints; with linear and quadratic terms), and their interaction (emotion X linear time), with a random intercept by subject. Emotion and time were modeled as repeated measures with a diagonal covariance structure and Satterthwaite estimates were used for determining degrees of freedom for the fixed effects. Separate models were fit for switch and stay conditions in the CO cluster (models for the L FP and R FP cluster are reported in the Supplemental Figure S2 and Tables S12-15).

### 2.6.5 Emotional Expression HRF Peak by Response Frame

Finally, to explore the effects of *response frame* on HRF timing and shape, a general linear model was fit according to the frame during which an individual participant made their judgment for each trial, as well as the emotional expression and response. Thus, there were regressors for positive happy, negative angry, negative fearful, positive surprised, and negative surprised trials for each response frame 2-5. Again, the “TENT” function was used to estimate the amplitude of the HRF at each TR from 0-21 seconds after stimulus onset. Regressors of no interest included error trials (negative happy, positive angry, and positive fearful responses), anticipatory responses (frame 1), polynomials for each run (two terms) and six motion parameters estimated during alignment (x, y, z shift/rotation). It should be noted that participants made most behavioral responses in frames 3 and 4 (∼40% each) and fewer responses in frames 2 and 5 (∼10% each), potentially impacting the stability of the HRF curve estimates across frames. All participants’ data were included for any condition in which a behavioral response was made. Beta values were extracted for each of the 16 ROIs and combined into the three ROI clusters of interest, focusing on the CO cluster (results for the L FP and R FP clusters are reported in the Supplemental Material). The average HRF for each emotion and frame was then upsampled from 22 time points to 2,200 using linear interpolation to aid visualization and minimize any impact of outlying time points, and peaks were identified using the “findpeaks” function in MATLAB. Peaks are reported that occurred at least five seconds into the trial (to allow the HRF time to develop), had an amplitude greater than 0.05 (to focus on positive activations), and were at least three seconds from another peak (to identify temporally distinct events).

Data are available online at https://osf.io/g39ke/.

## 3. Results

### 3.1 FMRI Study Behavior

Responses for valence judgments from the noise and reveal phases of the task were averaged separately for each of the four emotional expressions, resulting in a mean percent negative judgment (**Table 1**). Results from a linear mixed model indicated significant effects of emotion (*F*(3, 69.69) = 3170.72, *p* < .001), phase (*F*(1, 158.64) = 19.21, *p* < .001), and their interaction (*F*(3, 69.69) = 32.11, *p* < .001). As expected, happy faces were judged as mostly positive while angry and fearful faces were judged as mostly negative. Additionally, clear valence expressions showed a significant difference in percent negative judgments between the noise and reveal phase (*t*’s > 4.2, *p*’s < .001). As predicted, there were fewer errors (e.g., a happy face judged as negative) made during the reveal phase than the noise phase for each of the clearly valenced expressions. Judgments for surprised faces, on the other hand, did not significantly differ between the noise and reveal phases (*t*(31) = 0.87, *p* > .05) and were judged as somewhat negative during both phases, with more variability (i.e., individual valence bias) during the reveal phase than for the clear valence emotions.

**Table 1.**
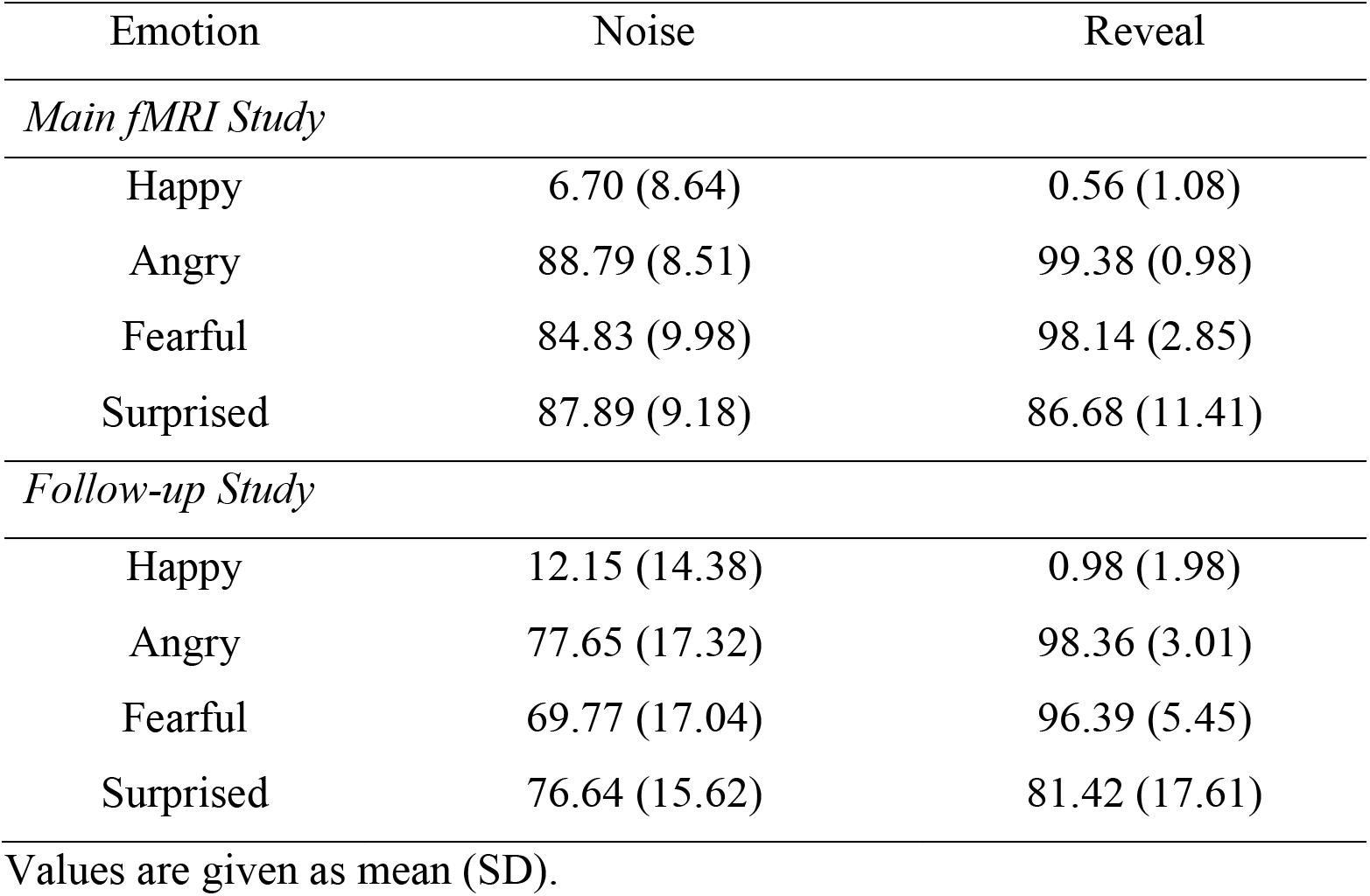
Average Percent Negative Judgments by Emotion.

| Emotion | Noise | Reveal |
| --- | --- | --- |
| *Main fMRI Study* |  |  |
| Happy | 6.70 (8.64) | 0.56 (1.08) |
| Angry | 88.79 (8.51) | 99.38 (0.98) |
| Fearful | 84.83 (9.98) | 98.14 (2.85) |
| Surprised | 87.89 (9.18) | 86.68 (11.41) |
| *Follow-up Study* |  |  |
| Happy | 12.15 (14.38) | 0.98 (1.98) |
| Angry | 77.65 (17.32) | 98.36 (3.01) |
| Fearful | 69.77 (17.04) | 96.39 (5.45) |
| Surprised | 76.64 (15.62) | 81.42 (17.61) |

Values are given as mean (SD).

Next, the noise frame (1-5) during which responses occurred was compared for correct happy, angry, and fearful face trials as well as separately for surprised face trials that were judged as positive versus negative. Descriptively, most responses for all emotions occurred during frames 3 and 4; happy faces that were correctly judged as positive and surprised faces that were judged as negative tended to have more early responses (frame 3) compared to correctly judged angry and fearful faces which had later responses (frame 4), as can be seen in the average response frame in the noise phase (**Table 2**). For a statistical test of these effects, a linear mixed model analysis was conducted on reaction times (RTs) for the noise phase and resulted in a main effect of emotion (*F*(4, 47.11) = 30.82, *p* < .001). Specifically, participants made slower responses for angry and fearful face trials and faster responses for happy and negative surprised face trials. For the reveal phase of the task, however, RTs showed a different pattern by emotion (*F*(4, 43.01) = 54.43, *p* < .001), with positive surprised trials resulting in the slowest responses, while happy trial responses were still faster than all other emotions (**Tables 2/S3**).

**Table 2.** Average Response Time by Emotion.

| Emotion | Average Frame | Noise RT (ms) | Reveal RT (ms) |
| --- | --- | --- | --- |
| *Main fMRI Study* |  |  |  |
| Happy | 3.25 (0.44) | 5417 (868) | 625 (118) |
| Angry | 3.61 (0.45) | 6124 (882) | 697 (120) |
| Fearful | 3.50 (0.48) | 5918 (949) | 699 (137) |
| Positive Surprised | 3.40 (0.69) | 5792 (1376) | 1018 (216) |
| Negative Surprised | 3.31 (0.51) | 5559 (1019) | 746 (168) |
| *Follow-up Study* |  |  |  |
| Happy | 4.82 (0.79) | 8515 (1588) | 742 (175) |
| Angry | 5.15 (0.74) | 9219 (1522) | 791 (202) |
| Fearful | 4.96 (0.72) | 8871 (1453) | 824 (191) |
| Positive Surprised | 4.82 (0.95) | 8695 (1917) | 1079 (259) |
| Negative Surprised | 4.81 (0.75) | 8637 (1481) | 831 (204) |

Values are given as mean (SD). The main fMRI study included five noise frames and the follow-up study included seven noise frames.

### 3.2 FMRI Data

#### 3.2.1 HRF Time Courses by Emotional Expression

The fMRI analysis of the HRF responses in the three ROI clusters of interest (as well as the Sensory control cluster) focused on differences as a function of emotional expression. For each ROI cluster separately, a linear mixed model was fit with fixed effects of emotion (5 levels), time (11 timepoints; with linear and quadratic terms) and their interaction (emotion X linear time), with reveal RT as a covariate and a random intercept by subject. Time points were included from the second half of the trial (11-21 seconds) when the BOLD response peaked, and emotions included clear correct trials (positive happy, negative angry, negative fearful) and both positive and negative surprised trials. See Supplemental Tables S5-S8 for full model results.

For the CO cluster, there were significant effects of emotion (*F*(4, 306.27) = 7.92, *p* < .001), time point (quadratic) (*F*(1, 696.42) = 49.37, *p* < .001), RT (*F*(1, 925.87) = 65.66, *p* <.001), and the interaction of emotion and time (linear) (*F*(4, 372.26) = 19.57, *p* < .001), and a non-significant effect of time point (linear) (*F*(1, 679.04) = 3.35, *p* = .068). The emotion effect and the interaction were driven by a stronger response for positive surprised trials from 13-21 seconds. Negative surprised trials also had a larger response than happy, angry, and fearful trials from 16 seconds until the end of the trial (**Figure 3A**).

For the L FP cluster, there were significant effects of emotion (*F*(4, 300.36) = 3.63, *p* =.007), time point (linear) (*F*(1, 640.71) = 17.14, *p* < .001), time point (quadratic) (*F*(1, 713.64) = 77.13, *p* < .001), RT (*F*(1, 904.89) = 16.90, *p* < .001) and the interaction of emotion and time (linear) (*F*(4, 350.84) = 5.10, *p* < .001). Similar to the CO cluster, positive surprised trials had a stronger response than the other emotions from 15-21 seconds (**Figure 3B**).

For the R FP cluster, there were significant effects of emotion (F(4, 281.99) = 15.62, p <.001), time point (linear) (F(1, 630.80) = 431.91, p < .001), time point (quadratic) (F(1, 716.23) = 507.72, *p* < .001), RT (*F*(1, 862.81) = 57.92, *p* < .001), and the interaction of emotion and time (linear) (F(4, 353.78) = 11.91, p < .001). The interaction was again driven by a stronger, more extended response for positive surprised trials compared to all other emotions beginning at 15 seconds (**Figure 3C**).

As a control, responses from five visual ROIs from Ploran et al. (2007) were averaged together in a “Sensory” ROI cluster. This cluster showed significant effects of time point (linear) (F(1, 594.89) = 742.64, p < .001), time point (quadratic) (F(1, 723.89) = 944.14, *p* < .001), and RT (*F*(1, 681.56) = 7.85, *p* = .005) with non-significant effects of emotion (F(4, 306.11) = 0.60, p = .660) and the interaction of emotion and time (linear) (F(4, 459.01) = 0.34, p = .850). All faces elicited a strong response in these regions that did not differ by emotional expression (**Figure 3D**).

#### 3.2.2 Late CO Activation and Behavior

Given the a priori interest in the CO network and evidence of a second, late peak for some conditions, the average BOLD activation was extracted for each emotion for the period from 16-19 seconds, which corresponds to 4-7 seconds after the fully revealed image was presented. Accordingly, we predicted that activation in this period would relate to behavioral responses for the reveal phase. A one-way ANOVA indicated that positive surprised trials had the strongest activation in this window (**Figure 4**), followed by negative surprised, while the clear valence correct trials (happy, angry, fearful) showed a relatively weak response (*F*(3.24, 67.95) = 37.07, *p* < .001, *η*^2^ = .638). Clear valence error trial activation was also extracted and did not significantly differ from positive (Bonferroni-corrected *p* = 1.00) or negative (*p* = .064) surprised trial activation.

**Figure 4.**
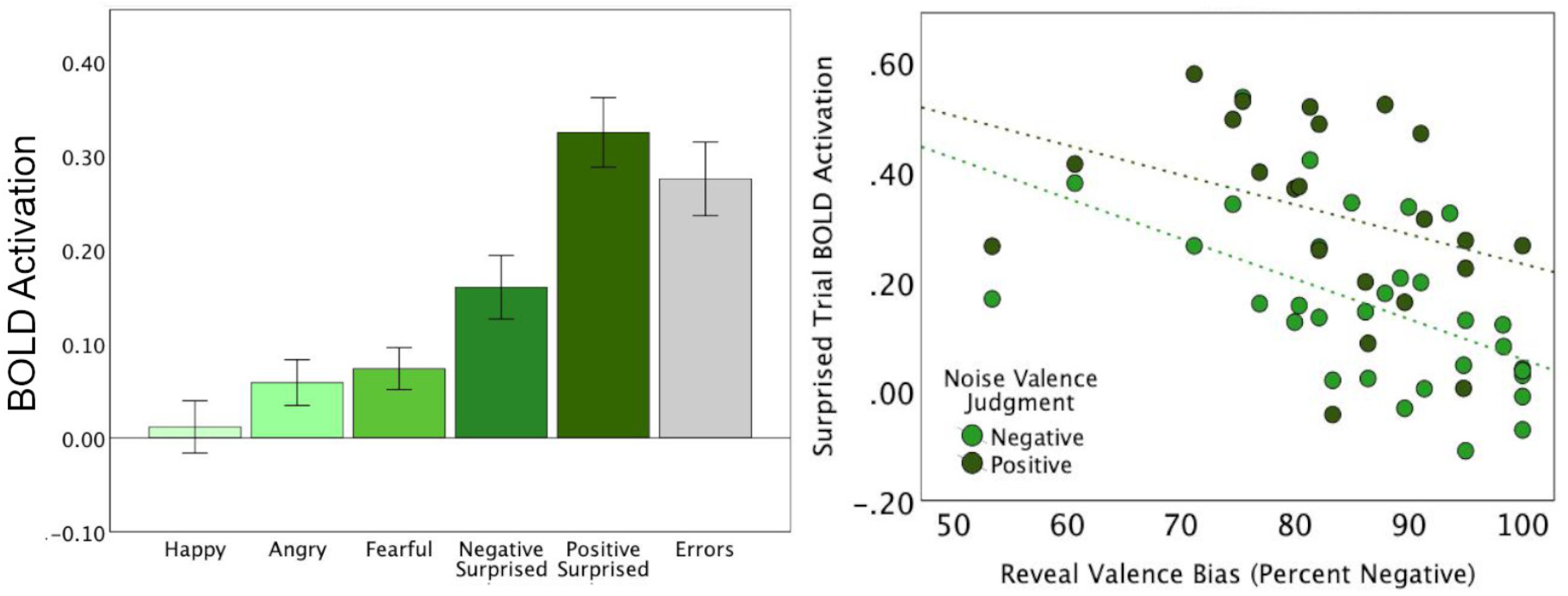
BOLD activation in the CO Cluster during a late trial window. *Left*: Average activation for each emotional expression during the late window (16-19 sec; 4-7 sec after reveal image), including clear valence error trials. *Right*: Correlation between late CO activation during surprised trials and the average valence bias (percent of negative judgments for surprised trials) during the reveal phase. For both negative and positive judgments during noise, participants with a more positive valence bias overall exhibited greater CO activation.

The correlation between each emotion’s late peak CO activation and its reveal RT was significant for fearful (*ρ* = .432, *p* = .014, 95% CI [.088, .684]) and negative surprised trials (*ρ* =.400, *p* = .023, 95% CI [.049, .663]), marginal for happy (*ρ* = .315, *p* = .079, 95% CI [-.048, .605]), and non-significant for angry (*ρ* = .206, *p* = .258, 95% CI [-.164, .525]) and positive surprised trials (*ρ* = .132, *p* = .567, 95% CI [-.330, .543]). This suggests that slower RTs may have contributed to some increased CO activation, but were not driving the marked increase for positive surprised trials. Additionally, the correlation between late peak CO activation and reveal valence bias was significant for both positive surprised (*ρ* = -.471, *p* = .027, 95% CI [-.750, - .048]) and negative surprised trials (*ρ* = -.635, *p* < .001, 95% CI [-.809, -.359]), indicating that individuals who made more positive judgments during the reveal phase had greater activation in the CO network regardless of their response for a given trial during the noise phase. (One participant was identified as a potential outlier based on reveal valence bias – removing this participant from the analysis did not significantly change the correlation values.) This finding may suggest that individuals with a more positive valence bias have a greater tendency to engage in CO-mediated regulatory processing of all ambiguous stimuli, even if they ultimately judge a specific face to have negative valence on a given trial.

#### 3.2.3 HRF Time Courses as a Function of Switching versus Staying

To explore in greater detail the nature of the HRF time courses for each emotional expression and assess whether error recognition (objective or subjective) at the full reveal was driving activation, each participant’s trials were sorted further according to whether they made the same response during the noise and reveal phases of the trial (i.e., “stay”) or switched to the opposite response, putatively indicating some type of error evaluation occurred (i.e., “switch”). As with the overall responses by emotional expression, a linear mixed model was fit with fixed effects of emotion, time (linear and quadratic), and their interaction (emotion X linear time), and a random effect of subject (intercept). Again, we focus on the CO cluster (**Figure 5**) given this network was previously associated with post-trial performance reporting (Gratton et al., 2017), including in response to ambiguity (Neta et al., 2017; Poudel et al., 2020), and error processing (Hester et al., 2004; Iannaccone et al., 2015; Neta et al., 2014), but the results from other clusters are shown in Supplemental Figure S2 and Tables S12-S15.

**Figure 5.**
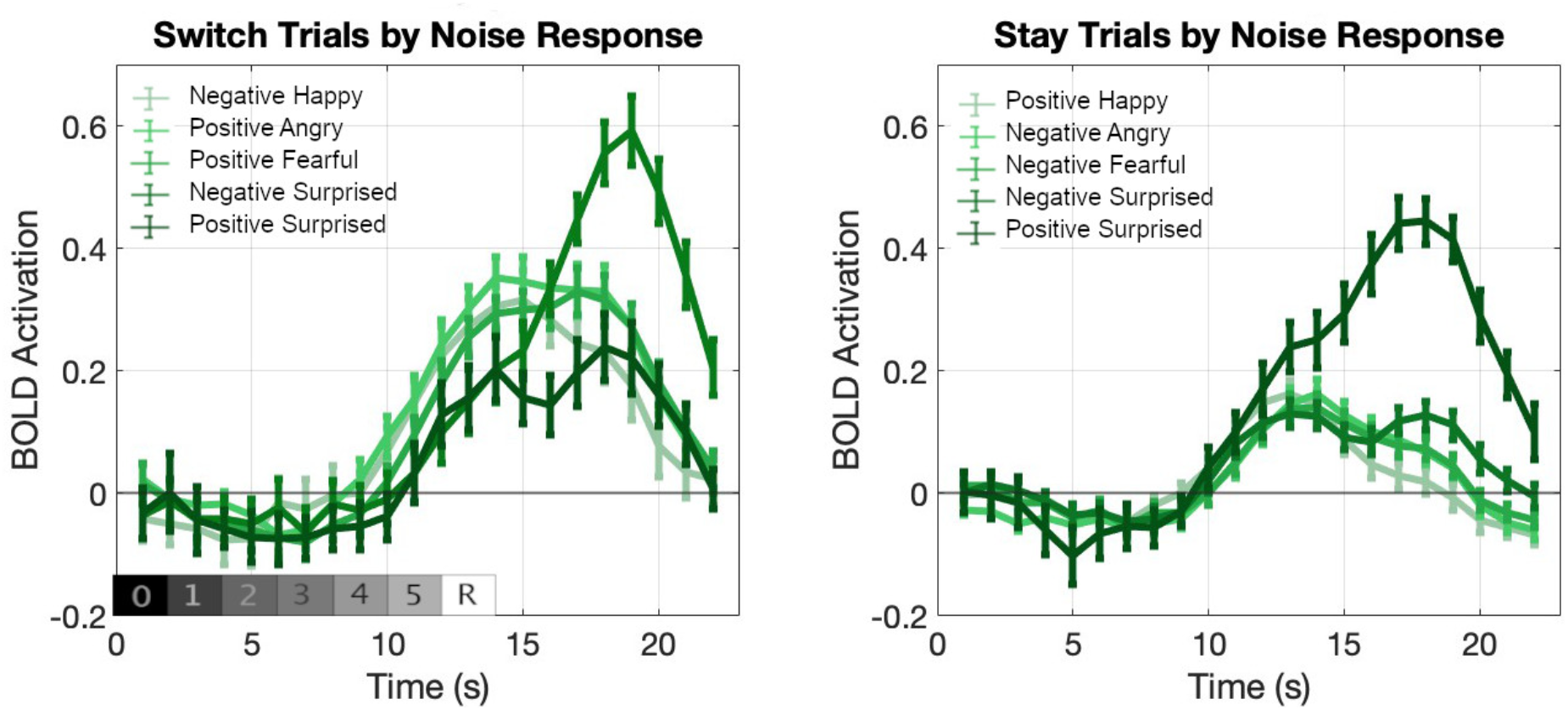
Estimated HRF curves in the CO ROI Cluster for each emotion based on whether a participant switched their behavioral response from the noise to the reveal phase or stayed with the same response. Labels refer to the initial response made during the noise phase. The gray bar represents the timing of each frame of the trial from 0 = black screen to 5 = final noise frame and R = fully revealed image.

For switch trials in the CO cluster, there were significant effects of emotion (F(4, 199.82) = 26.09, p < .001), time point (linear) (F(1, 476.73) = 181.64, p < .001), time point (quadratic) (F(1, 510.66) = 240.32, *p* < .001), and the interaction of emotion and time (linear) (F(4, 204.77) = 45.33, p < .001; see Supplemental Table S10). Descriptively, surprised trials for which participants responded “positive” during the noise phase showed the weakest response, while clear valence error trials (negative happy, positive angry/fearful) had a stronger response. Critically, surprised trials for which participants responded “negative” during noise and “positive” at reveal exhibited a moderate response from 11 to 14 seconds, but showed the strongest response from 16 seconds to the end of the trial (i.e., after the image had been revealed and the positive judgment made).

For stay trials, there were significant effects of emotion (F(4, 220.56) = 6.79, p < .001), time point (quadratic) (F(1, 684.90) = 42.72, *p* < .001), and the interaction of emotion and time (linear) (F(4, 281.60) = 16.21, p < .001), and a non-significant effect of time point (linear) (F(1, 687.95) = 1.86, p = .173; see Supplemental Table S11). Correct clear valence trials (positive happy, negative angry/fearful) and negative surprised trials showed a relatively weak response. Importantly, positive surprised trials showed the strongest response, particularly after 14 seconds, consistent with the switch trial results in demonstrating that the largest response occurred when the final valence judgment of surprised faces was positive.

#### 3.2.4 Emotional Expression HRF Peak by Response Frame

Finally, the HRFs for each emotional expression were analyzed according to the frame during which a behavioral response was made to examine how the timing of the noise judgment impacted the HRF timing and shape. In the CO cluster during clear valence trials, the HRF generally had a similar, standard shape with a shift in peak time relative to the timing of the noise behavioral response (**Figure 6**; see Supplemental Table S16 for exact peak times). For surprised trials, however, the HRF time courses for early frames showed two peaks: the first which corresponded to the clear valence trial peaks and the second which occurred after the reveal image was shown. This double-peak effect may indicate that, specifically for surprised trials, the participants processed the valence of the face not only during the noise phase, but perhaps also completed some additional processing or performance reporting after the full image was revealed. This second level of processing is likely attributed to the ambiguity of the valence of the surprised expression. (The responses by frame for the other ROI clusters are shown in Supplemental Figure S3.)

**Figure 6.**
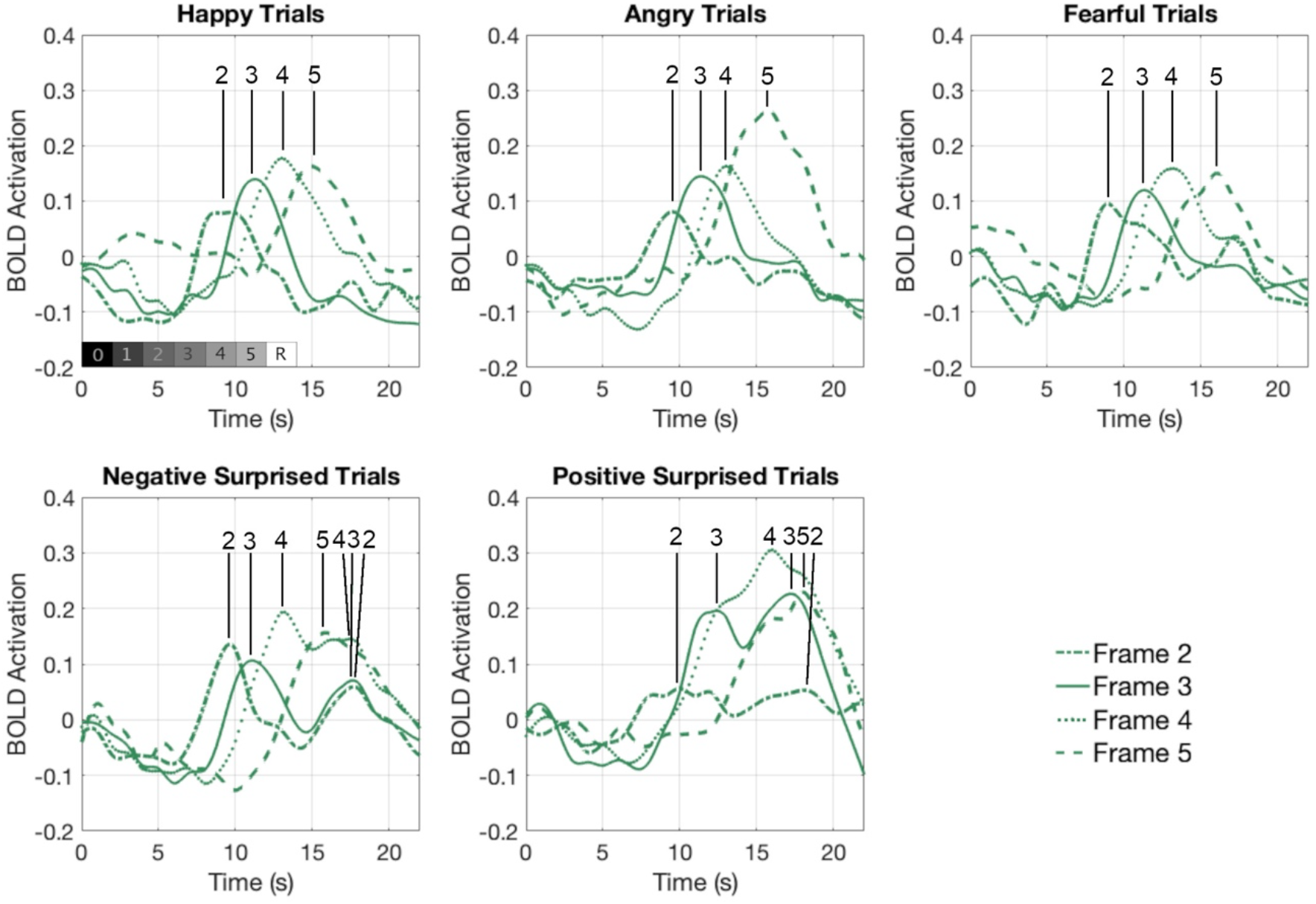
Estimated HRF curves for each emotional expression by response frame for the CO cluster. Averaged responses across participants according to the noise frame (2-5) during which a behavioral response was made for correct clear valence trials (happy, angry, fearful) and ambiguous valence (surprised) trials for negative and positive responses. Numbers at the top of each plot indicate the identified peak(s) for each frame.

#### 3.3 Follow-up Online Study

In order to further tease apart the behavioral response pattern observed in the main study, particularly for surprised faces that are judged to have positive versus negative valence, a follow-up study was conducted online. Given that most responses were made during noise frames 3 and 4 in the main study and there was thus a limited window to distinguish the timing of ambiguity judgments, the slow reveal paradigm here was modified to include seven (instead of five) frames of each facial expression in the “noise” phase of the trial. This was achieved by creating smaller steps between frames such that the perceptual information was revealed more slowly, rather than extending the noise frames towards a more fully revealed image (a full description of the methods is provided in the Supplemental Material). The longer period during which the face was unmasked was intended to allow for greater sensitivity to perceptual changes that lead to variability in response frame. With this extended window for noise responses and based on the initial negativity hypothesis, it was predicted that for surprised faces positive judgments would be made later than negative judgments during the noise phase as well as the reveal phase.

The final sample consisted of 81 naïve participants (39 F/42 M) with a mean age of 31.1 years (range 18-39). Responses for valence judgments from the noise and reveal phases of the task were averaged separately for each of the four emotional expressions, resulting in an average percent negative judgment (**Table 1**). Results from a linear mixed model indicated significant effects of emotion (*F*(3, 192.23) = 1962.40, *p* < .001), phase (*F*(1, 406.77) = 97.53, *p* < .001), and their interaction (*F*(3, 192.23) = 91.00, *p* < .001). Consistent with the results from the main study, happy faces were judged as mostly positive while angry and fearful faces were judged as mostly negative, with fewer errors (e.g., a happy face judged as negative) made during the reveal phase than the noise phase for each of the clearly valenced expressions (*t*’s > 7.10, *p*’s < .001).

Unlike the main study, surprised faces were judged as significantly more negative during the reveal phase than the noise phase (*t*(80) = 3.59, *p* < .001, Cohen’s *d* = .399), albeit with a smaller difference between phases than the clear valence emotions. Next, the frame (1-7) during which responses occurred during the noise phase was compared for correct happy, angry, and fearful trials as well as separately for surprised trials that were judged as positive and negative. Descriptively, most responses occurred during frame 5 for all emotions (**Table 2**), but happy faces that were correctly judged as positive and surprised faces that were judged as negative yielded earlier responses (frames 3 and 4) than angry and fearful faces that were correctly judged as negative (frames 6 and 7). A linear mixed model analysis with a fixed effect of emotion on RT during the noise phase resulted in a main effect of emotion (*F*(4, 113.62) = 27.05, *p* < .001). Participants made slower responses for angry and fearful face trials and faster responses for happy and negative surprised face trials. As in the main study, the reveal phase of the task showed a different RT pattern by emotion (*F*(4, 108.29) = 48.39, *p* < .001), with positive surprised trials resulting in the slowest responses, while happy trial responses were faster than all other emotions (**Tables 2/S4**).

Surprised trials for which participants stayed with the same response for the reveal phase were subsequently analyzed according to the percentage of responses in each frame (given that some participants did not make any positive stay responses, the sample size was reduced to 68 for this analysis). There was a significant effect of frame (*F*(2.43, 279.70) = 34.79, *p* < .001, *η*^2^ = .342) and a significant interaction between frame and the valence of the response (*F*(4.09, 273.82) = 3.92, *p* = .004, *η*^2^ = .055). Positive surprised stay trials were rated later than negative surprised stay trials. Specifically, pairwise comparisons indicated that during frames 3 and 4 more negative responses were made, while during frame 5 more positive responses were made (**Figure 7**). This effect was marginal for RT (stay positive: 9026 (2230) ms; stay negative: 8672 (1553) ms. *t*(67)=1.53, *p*=.066, Cohen’s *d*=.185).

**Figure 7.**
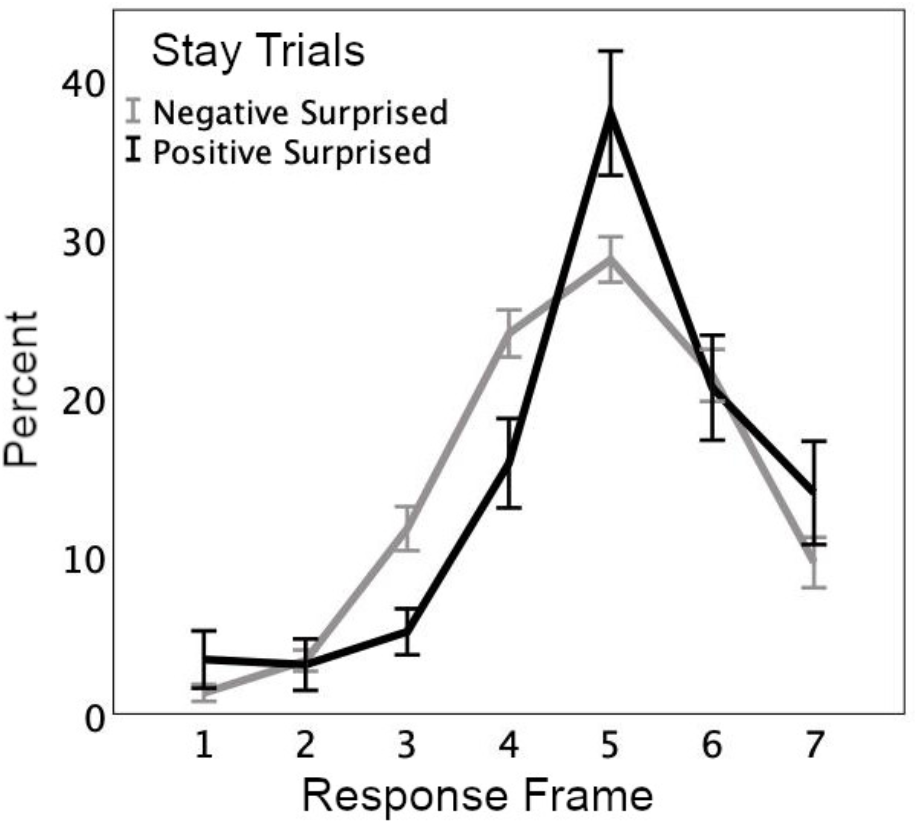
Percent of responses during each noise frame in the follow-up study for trials on which participants stayed with the same response for the reveal phase.

## 4. Discussion

In the current work, a valence bias task was presented in which clearly and ambiguously valenced emotional facial expressions were slowly revealed over several seconds and participants were asked to make a valence judgment (positive/negative) about each face both during the unmasking and after the full image was presented. This design allowed for examination of the decision-making mechanisms involved in processing and resolving ambiguity as they unfold over the course of a trial. Overall, ambiguous (surprised) trials on which the participant judged the face to have positive valence exhibited the most unique response characteristics and strongest activation in the CO network late in the trial, supporting the initial negativity hypothesis of ambiguity processing.

In the main study, BOLD HRF time courses were extracted from 16 ROIs in three cortical networks previously associated with task control, decision making, and performance reporting. The results demonstrated greater activation for ambiguous (surprised) trials in the CO network – and more so for positive judgments of surprised faces – particularly in the latter part of the trial, which included a second activation peak following the presentation of the reveal image. Moreover, the CO activation for surprised faces correlated with valence bias, such that individuals with a more positive bias showed increased activation in the late stages of the trial. This pattern of increased activation including a second peak at the final decision on ambiguous trials is in line with a performance reporting role of this network.

In a follow-up study, the task was administered to a larger, online sample using a more extended slow reveal design with additional noise frames. Across both studies, participants responded more slowly when judging surprised faces as positive versus negative. Taken together, these findings support the initial negativity hypothesis (Neta, 2024; Neta & Whalen, 2010), which posits that the default response to ambiguity is negative and that a positive evaluation requires greater effortful control or regulation to overcome this initial negativity. Specifically, these findings indicate that this regulatory process is supported by CO network activity.

### 4.1 Unfolding of Valence Judgments Differs by Emotion

Behaviorally, on clear valence trials participants made more errors (e.g., judging a happy face to be negative) during the noise phase when the face was partially obscured, which they subsequently switched to correct responses during the reveal phase when the full image was visible. This is consistent with the fact that the masked expressions during the noise phase were less perceptually clear and participants may have confused what they saw with an expression signaling the opposite valence. Conversely, surprised trials showed no difference in average valence judgments between the two task phases in the fMRI study, suggesting that participants evaluated the surprised faces during the reveal in accordance with their initial response during the noise phase. In other words, the perceptual uncertainty of the slow reveal paradigm may not have influenced their judgments strongly and participants responded instead according to their own internal valence bias.

Participants in the online study, however, judged surprised faces as more positive during the noise phase than during the reveal phase. The elevated number of positive judgments (i.e., lower percent negative) for surprised faces may indicate that more participants in this larger sample made more subjective errors (e.g., a positive judgment that they later switched to a negative judgment) due to the perceptual uncertainty during the noise phase – as was the case for clear valence trials. An alternate explanation could be that the extended slow task design allowed participants time to adopt a positive interpretation of the surprised face during noise (see Neta & Tong, 2016), that was nonetheless overturned during the subsequent quick judgment at reveal.

Another possibility is that the extended viewing of a face through the full trial allowed participants to accumulate sufficient perceptual evidence to make a “correct” negative judgment of surprise at reveal. Unfortunately, the current design does not allow for investigation of participants’ motivation when switching their response, so future work will be needed to better understand this pattern of results.

Response times exhibited no significant difference between surprised faces judged as positive versus negative during the noise phase in the fMRI study or in the initial analysis in the follow-up study. However, given that participants occasionally switched their valence judgment from the noise to the reveal phase for surprised trials (possibly indicating a subjective or motoric error), we then compared response times for surprised trials on which a participant stayed with the same response during both phases in the follow-up online study. For these “stay” surprised trials, more negative responses were made in an earlier frame and more positive responses were made in a later frame during the noise phase. This result mirrors the surprised RT effects during the reveal phase and supports the idea that positive judgments of ambiguity require longer processing time to overcome an initial negativity, consistent with prior work (Neta & Tong, 2016; Petro et al., 2021), even in a context with perceptual uncertainty.

### 4.2 Disentangling Perceptual and Valence Ambiguity

Performance in the current task required two valence judgments about an emotional face that was slowly unmasked and then fully revealed. When considering the behavioral responses from both studies, we found evidence that distinct factors may contribute to decision-making processes during the noise and reveal phases of the task. During the noise phase, participants responded to the happy faces faster than the angry and fearful faces, indicating that perceptual features may be contributing to response times. Assuming that participants first recognized the expression or some subset of its physical features (Calvo & Nummenmaa, 2016) prior to making a decision about its valence (Storbeck et al., 2006), it may be that differences in the morphological features of the expressions impacted response times most strongly during the noise phase. In other words, happy faces are easiest to recognize, likely due to the unique appearance of a smile (Calvo et al., 2012). Relatedly, other emotional expressions that are perceptually similar (e.g., both surprise and fear have widened eyes) may need more time/information to recognize the expression and categorize its valence (Calvo & Lundqvist, 2008; Calvo & Nummenmaa, 2016; Tottenham et al., 2009), which likely explains the longer response times for fearful faces.

This is in contrast to the reveal phase, when participants still responded faster to happy than angry or fearful faces, but now responded the slowest to surprised faces that they judged as positive, as predicted. Thus, when the face was not obscured and there was no perceptual uncertainty, the morphological similarities had a weaker effect on the decision process than the valence ambiguity of surprise. This is consistent with prior work examining valence bias (Harp et al., 2021; Neta et al., 2009; Neta & Whalen, 2010) and with the decision-making literature more broadly, where ambiguous trials result in slower responses or response aversion (Budescu et al., 2002; Ellsberg, 1961; Ikink et al., 2019).

### 4.3 Task Control Networks Support Ambiguity Processing

The CO, R FP, and L FP control networks make variable contributions to decision-making and task control that can be distinguished in part by their temporal characteristics. Previous research has investigated how these networks accumulate evidence, handle errors, and report task performance during the gradual reveal of objects or words in identification tasks (Gratton et al., 2017; Neta et al., 2017; Ploran et al., 2007; Wheeler et al., 2008). When examining HRF time courses in response to emotional faces (see Supplemental Material), we replicated prior work characterizing temporal characteristics of task control networks using slow reveal tasks (Gratton et al., 2017). This replication supports the domain-general decision-making functions of these regions and demonstrated the temporal effects using a more complex and socially relevant stimulus set that required a distinct type of decision to be made (i.e., categorizing emotional valence).

Critically, when the HRF was examined in these networks according to the emotional expression of the face, there was a marked increase in BOLD activation for surprised trials, particularly when a participant judged the face’s valence as positive, as compared to trials with clearly valenced expressions. The surprised trial response was most notable in the CO network, with a strong response late in the trial for positive judgments and to a lesser extent negative judgments as well. In contrast, a sensory cluster composed of early visual regions showed no difference in activation by emotion and was engaged earlier in the trial as sensory information became available, well before a valence decision was made.

Further examination of the ambiguity-related late BOLD response in the CO network indicated a relationship with individual behavior such that greater activation during the noise phase of surprised trials was associated with a more positive valence bias (i.e., one’s tendency to judge the expressions as positive). Therefore, individuals who have a more positive valence bias may process the ambiguous trials more thoroughly or effortfully regardless of their ultimate decision on a specific trial. This is consistent with other work demonstrating that physiological responses to emotional ambiguity track one’s general tendencies (i.e., valence bias) rather than their transient response on a given trial (Neta et al., 2009). Additionally, an examination of the clear valence error trial activation (combined across happy, angry, and fearful) demonstrated a moderate response that did not significantly differ from surprised trials. This is consistent with previous work showing that ambiguity, RT, and error commission represent separable responses within the CO network (Neta et al., 2014).

### 4.4 Ambiguity Elicits a Second BOLD Peak following the Second Valence Judgment

To investigate how the HRFs differed as a function of *when* the first valence decision occurred, activation in the CO network was analyzed according to the frame during which a participant responded. Clear valence expression trials showed a single HRF peak that was shifted in time relative to the response frame. Surprised trials showed a similar first peak, yet unexpectedly also exhibited a second HRF peak late in the trial. This peak was most evident for responses in frames 2 and 3, whereas for responses in frames 4 and 5, the two peaks seemed to overlap temporally. Similarly, the two peaks were less distinct in the prior analyses combined across frames, but nonetheless are visually apparent in the CO surprised trial HRFs (Fig. 3A).

The late peak presumably corresponds to a second judgment following the full reveal of the face image, which evidently was not necessary for clear valence expressions. Interestingly, this second peak occurred for both positive and negative surprised trials, indicating that this emotional expression not only modulated the strength of the CO activation, but also the shape of the HRF, as a result of distinct decision-making demands for ambiguous stimuli. Prior work using the slow reveal paradigm did not exhibit such a pattern for object recognition (Gratton et al., 2017; Neta et al., 2017), likely because once the image was fully revealed, the perceptual ambiguity was resolved. In other words, a picture of a toothbrush might look like many things when only partial visual information is available, but once the full image is presented, the object is clearly identifiable. In contrast, in the current work, the full reveal of an ambiguously valenced (surprised) face is not sufficient for ambiguity resolution, so it elicits a second peak that may indicate participants were not simply confirming their first judgment, but perhaps making a distinct second decision about its valence. Nonetheless, the number of responses for each expression and frame was fairly small, so these results must be considered preliminary.

Although this study was not designed specifically to test the CO’s performance monitoring role, the present findings regarding the CO network’s response to ambiguity support previous evidence of its functional role in performance reporting following a task decision (Gratton et al., 2017, 2018; Neta et al., 2014, 2017; Ploran et al., 2007). Activation in this network began to increase later than sensory or FP ROIs and as a function of the behavioral response timing. Speculatively, this late activation is consistent with a performance reporting role that occurs once a (valence) decision has been made, with differing response strength based on the judgment that was selected. Specifically, a positive judgment of surprise evoked a stronger late CO response than a negative judgment did, indicating that the outcome of the decision-making process was relevant to this network’s control function. Prior work has shown that activation in the CO network during ambiguity led to improved accuracy on subsequent trials (Neta et al., 2017), demonstrating that its performance reporting role can impact ongoing behavior.

Alternate explanations of the function of CO regions, including salience detection, task set maintenance, conflict monitoring, and alertness (Botvinick et al., 2001; Coste & Kleinschmidt, 2016; Dignath et al., 2020; Dosenbach et al., 2008; Han et al., 2019; Seeley et al., 2007), do not align as well with our results given the specific *late* increase in activation for ambiguous over other emotional faces. It could be argued, however, that ambiguity might signal increased trial conflict or difficulty, carry greater affective salience, and/or necessitate heightened attention. Nonetheless, the late onset of the CO response (i.e., after a decision has been made) supports a performance reporting role that is engaged at the end of an evaluation rather than an ongoing monitoring or attentional function (see also Neta et al., 2014).

### 4.5 Evidence in Support of the Initial Negativity Hypothesis

The increased activation for positive surprised trials in the CO network supports the proposal regarding the primacy of a negative judgment of ambiguity, with a positive judgment requiring additional top-down processing to overcome this initial assessment (Neta & Whalen, 2010; Petro et al., 2018). We examined two possible behavioral explanations for the unique effects for positive judgments of surprise. The first was reaction time, which had a significant effect in the HRF mixed models such that slower responses were associated with greater activation in each network. Nonetheless, the effect of emotional expression was significant even when accounting for RT. This finding was supported further by the correlations between reveal RTs and late CO activation, which were significant for fearful and negative surprised trials, but not for positive surprised trials. This pattern of results demonstrated that while processing time may contribute to CO activation, the large response for positive surprised trials was not merely an artefact of slower processing, but likely reflects a specific control process necessary to resolve the ambiguity.

Secondly, the increased activation in the CO network for positive surprised trials was analyzed according to whether a participant switched their response from the noise to reveal phase or stayed with the same response. We found that making a positive judgment of a surprised face during the reveal phase resulted in the greatest activation in the CO network regardless of the initial behavioral response (i.e., negative surprised switch trials and positive surprised stay trials had the largest late activation). This result implies that resolving the ambiguity of a surprised face with a positive judgment creates unique decision-making or regulatory demands, or perhaps entails repeated performance reporting following an uncertain initial decision. One alternative explanation is that a difference in difficulty between ambiguous and clear trials (Desender et al., 2021) contributed to the greater activation for positive surprised trials, yet this account would still be consistent with the initial negativity hypothesis.

The current results also indicated that the large response for positive surprised trials was not a type of error signal, as even those trials on which participants stayed with a positive judgment during the reveal phase showed increased activation. Furthermore, clear valence trials where participants made an erroneous response during the noise phase and switched their valence judgment to the correct response during the reveal phase showed only a moderate response that was initially stronger than the surprised trials but weaker than the negative surprised switch trials in the later part of the trial.

Collectively, the current results demonstrate clear support for the initial negativity hypothesis, which proposes that most people initially form a quick negative appraisal of an ambiguous image, perhaps in service to preparing a response to a potential threat. Then, according to the individual’s internal biases, they may reevaluate the stimulus to find a more positive appraisal (see also Neta et al., 2022). This process seems to be supported in part by the CO network, given the general increased activation for positive surprised trials and the relationship between a positive valence bias and the extent of this activation.

### 4.6 Limitations and Future Directions

The current results must be considered with respect to some limitations. First, positive judgments for surprised trials did not occur frequently and differed in frequency across participants, potentially impacting the nature of this response relative to other emotional expressions. This issue is further amplified in the exploratory analyses examining switch vs. stay trials and responses by frame, which divided positive surprised trials in an effort to investigate possible explanations for the unique responses. Further work with a greater number of trials or individuals with a more positive valence bias is needed to determine whether or how the infrequency of this condition impacts the HRF characteristics.

Second, the task design does not allow for separation of the HRFs for the noise and reveal phases, so it is not possible to precisely distinguish the contribution of each phase. This may impact the interpretation of the secondary, late peak for ambiguous trials, which was unexpected. Yet it is reasonable to conclude based on its timing that the later peak is driven by the presentation of the fully revealed face. Future work could explore the response to distinct phases of ambiguity processing in greater detail. For example, a task design with a delay between the noise and reveal phases or catch trials during which no reveal image appeared could help illuminate the nature of the two apparent peaks.

Finally, the average surprised valence judgments during the reveal phase in both studies were relatively negative compared to prior reports (e.g., Harp et al., 2022; Neta et al., 2009; Petro et al., 2018). It may be that the greater number of clearly negative facial expression conditions (angry and fearful vs. happy) created a negative context (Neta et al., 2011) that biased participants towards making a negative response or that the specific surprised faces selected for this study tend to elicit more negative judgments. Importantly, however, surprised faces were immediately preceded by an equal number of clearly positive and negative faces. This point warrants further investigation to determine whether this negativity is a result of a negatively biased context or if perhaps the nature of the slow reveal paradigm itself somehow shifts valence bias due to increased perceptual uncertainty or slow pacing.

### 4.7 Conclusion

Ambiguous stimuli often occur in daily life within contexts that may not provide sufficient evidence for a single clear interpretation. In these two studies, we took a novel approach using a perceptual decision-making task to explore the mechanisms that support the unfolding of judgments about ambiguous and clear facial expressions. We demonstrated that cortical task control networks support processing of emotional facial expressions as they are slowly revealed. Positive judgments of surprised faces were accompanied by stronger activation in the CO network and slower response times, supporting the initial negativity hypothesis. Surprised faces also resulted in a second activation peak in the HRF following the full reveal, which likely reflects the continued valence ambiguity of these stimuli. Together, these findings point to a role of the CO network in late responses to ambiguity, in line with a performance reporting role. Further, the regions in this network seem to support the effortful or regulatory process that particularly supports more positive judgments of ambiguity.

Funding: This publication was made possible by Nebraska Tobacco Settlement Biomedical Research Development Funds and the National Institutes of Health (NIMH111640; PI: Neta).

The authors declare that there are no competing financial interests.

## Supplemental Methods

### HRF Response Characteristics by ROI Cluster

The data from all runs were entered into a general linear model with regressors based on each individual participant’s noise responses (from frames 2-5), collapsing across all emotion expressions. The “TENT” function was used to estimate the amplitude of the hemodynamic response at each TR from 0-21 seconds after stimulus onset (22 timepoints; TR = 1 second) without assuming a predetermined shape for the response in each voxel. Regressors of no interest included anticipatory responses (frame 1), error trials (incorrect judgment for a clearly valenced stimulus), polynomials for each run (two terms), and six motion parameters estimated during alignment (x, y, z shift/rotation).

To compare the response timing and shape in each of the three ROI clusters of interest, several parameters characterizing the HRF were extracted. Following the approach of Gratton et al. (2017), peak time, onset time, offset time, and full width-half maximum (FWHM) were quantified for each ROI cluster in each participant. First, time courses from each ROI were averaged to create an estimated HRF for each ROI cluster for each participant. Next, these average HRFs were expanded from 22 time points to 2,200 time points using linear interpolation in MATLAB to obtain better characterization of the underlying HRF and limit the influence of any potential outlying single data points. From these interpolated time courses, peak time was defined as the time point at which the curve reached its maximum amplitude. Onset time was defined as the time point at which the curve exceeded 25% of the peak amplitude (calculated by stepping back from the peak), while offset time was defined as the time point at which the curve dropped below 25% of the peak amplitude after the peak time. FWHM was defined as the time between the points at which the curve rose above and fell below 50% of the peak amplitude. A repeated measures ANOVA was conducted to compare the response characteristics between ROI cluster (3 levels) for peak time, onset, offset, FWHM, and peak amplitude. Huynh-Feldt adjusted degrees of freedom were used to account for unequal variance between conditions when indicated by a significant Mauchly’s test of sphericity and Bonferroni corrected *p*-values were used for pairwise comparisons.

## Follow-up Online Study

### Participants

Data was collected from 135 participants recruited online using Gorilla Experiment Builder (Anwyl-Irvine et al., 2020). Participants were required to be at least 18 years old and residing in the United States. Participants’ data were excluded if at least half of the task was not completed, participants did not respond to attention checks (press ‘A’ or ‘L’ when cued), or if clear valence expressions were not judged correctly at the reveal (e.g., judging most happy faces to be negative). This resulted in a final sample of 81 participants (39 F/42 M) with a mean age of 31.1 years (range 18-39), and race/ethnicity reported as 75% White, 10% Asian, 5% Black, and 5% Hispanic/Latino. Participants received monetary compensation for their time. All task materials and procedures were approved by the UNL Institutional Review Board.

### Task Design and Procedure

Each trial began with the presentation of a white fixation cross in the center of a black screen for 250 ms, followed by a black screen for 2 seconds. Then, a grayscale face was presented that was occluded by a black mask that was slowly degraded over the course of seven 2-second frames. The black mask was degraded by randomly removing pixels following a Gaussian distribution over the seven “noise” frames such that 3.5, 5, 6.5, 8.5, 11, 13.5, and 16% of the face was visible at each successive step. The trial ended with the presentation of the fully revealed face stimulus for 2 seconds. Thus, the duration of each trial was 18 seconds from the onset of the black screen to the offset of the face stimulus.

As in the main study, participants were instructed to judge each face as either positive or negative twice during the trial, once when the picture was emerging and again when the picture was fully revealed, and were told that the two responses could be the same or different. The participants completed ten practice trials containing only angry and happy faces as examples of negative and positive valence (no surprised faces) that were not used in the main experiment, for which they received feedback as to whether they correctly identified the image’s valence during the noise and reveal stages. There was a total of 120 trials (30 unique images each of happy/angry/fearful/surprised), with faces selected from the NimStim Set of Facial Expressions and the Karolinska Directed Emotional Faces database. Participants used a keyboard button (“A” or “L”) to rate each image with the mapping to positive/negative counterbalanced across participants.

### Data Analysis

As in the main study, the average valence judgments for each emotion were calculated as the number of negative responses out of the number of total trials for each participant, multiplied by 100 (i.e., percent negative). Average judgments were entered into a linear mixed model with fixed effects of emotion (4 levels - happy, angry, fearful, surprised), trial phase (2 levels – noise, reveal), and their interaction, with a random intercept by subject. Emotion and phase were modeled as repeated measures with a diagonal covariance structure and Satterthwaite estimates were used for determining degrees of freedom for the fixed effects. Subsequently, incorrect judgments for clearly valenced emotional expressions (happy judged as negative and angry or fearful judged as positive) were considered errors and not included in the analyses.

Next, the average response frame (out of 7) was calculated for each of the five emotion conditions (happy, angry, fearful, positive surprised, and negative surprised) for each participant. The noise RT was calculated from the beginning of the first frame, giving a maximum possible value of 14 seconds (2 sec x 7 frames), while the reveal RT had a maximum possible value of 2 seconds. Noise and reveal RTs were entered into separate linear mixed models with fixed effects of emotion (5 levels) and a random intercept by subject. Emotion was modeled as a repeated measure with a diagonal covariance structure and Satterthwaite estimates were used for determining degrees of freedom for the fixed effects.

To further examine the behavioral relationship between positive and negative responses on surprised trials, the percent of responses made during each of the seven frames was compared depending on whether the participant stayed with the same response during the noise and reveal phases. Given the low number of “switch” surprised trials, only noise trials that were judged as having the same valence during the reveal phase were included. These “stay” surprised trials are presumably those for which the participant made a subjectively “correct” judgment during the noise phase and, thus, confirmed this judgment during the reveal phase. As some participants did not make any positive judgments of surprised faces during the reveal phase, this resulted in a reduction of the sample size to 68 for this analysis.

## Supplemental Results

### Response Characteristics by ROI Cluster

This analysis focused on differences in response characteristics of the estimated HRF curve across three ROI clusters (CO, L FP, R FP) created from 16 ROIs from Gratton et al. (2017). Based on the average HRF in each ROI cluster (**Figure S1**), onset time, offset time, peak time, peak amplitude, and FWHM were calculated for each participant and compared across ROI clusters using one-way ANOVAs. There was a significant effect of ROI cluster in all measures (Figure S1). For onset time, there was a significant effect of ROI cluster (*F*(1.75, 54.16) = 5.22, *p* = .011, *η*^2^ = .144), with the L FP cluster response rising earlier than the R FP response (Bonferroni corrected *p* = .026). For peak time (*F*(1.51, 46.76) = 10.00, *p* < .001, *η*^2^ = .244) and offset time, (*F*(2, 62) = 6.10, *p* = .004, *η*^2^ = .164), the pairwise comparisons indicated that the R FP response peaked later than L FP (*p* = .015) and ended later than both other regions (R FP vs L FP: *p* = .028; R FP vs CO: *p* = .011). For FWHM, there was also a significant effect (*F*(1.68, 52.17) = 6.29, *p* = .006, *η*^2^ = .169), such that the CO had a smaller FWHM than the R FP cluster (*p* < .001). Finally, there was a significant effect for peak amplitude (*F*(2, 62) = 3.88, *p* = .026, *η*^2^ = .111), such that the R FP response had a larger amplitude than the L FP response (*p* = .035). Across all emotions, the R FP network had a stronger, later, more extended response, while the L FP and CO networks had a more transient response, consistent with the findings of Gratton et al. (2017). The L FP network also became activated earlier and with a smaller amplitude than the other networks. These results imply that the control networks are performing similar decision making and performance reporting functions in the slow reveal paradigm regardless of the specific stimulus presented or task decision that must be made.

---

**Figure S1.**
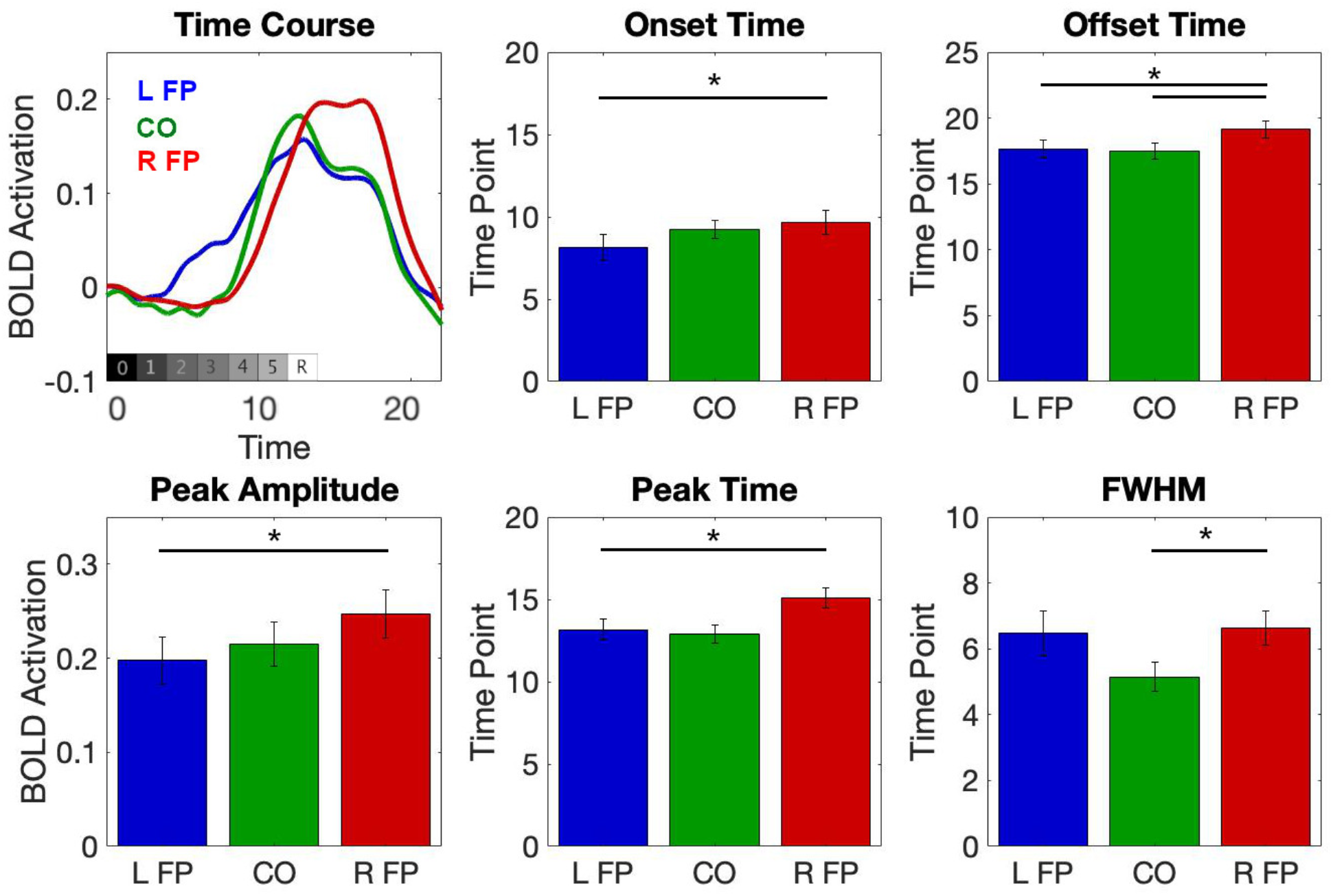
Estimated HRF curves of the three ROI clusters. The upper left panel shows the interpolated HRF curve averaged from each of the ROIs within the three network clusters across participants. The gray bar represents the timing of each frame of the trial from 0 = black screen to 5 = final noise frame and R = fully revealed image. The L FP cluster had the earliest onset and smallest amplitude, while the R FP cluster had a later onset, offset, and peak time with the largest amplitude, and the CO cluster had the smallest FWHM. The bar graphs illustrate the metrics derived from these curves to characterize differences in shape and timing across clusters.

**Table S1.** MNI Coordinates for each region of interest.

| Region | x | y | z |
| --- | --- | --- | --- |
| *Cingulo-opercular cluster* |  |  |  |
| Left AI/FO | 31 | -25 | -1 |
| Right AI/FO | -33 | -26 | 1 |
| Pre-SMA 1 | -1 | -15 | 51 |
| Pre-SMA 2 | -7 | -24 | 37 |
| *Left frontoparietal cluster* |  |  |  |
| Left IPS 1 | 42 | 41 | 45 |
| Left IPS 2 | 32 | 50 | 45 |
| Left IPS 3 | 24 | 67 | 49 |
| Left frontal | 43 | -4 | -33 |
| Left MFG 1 | 45 | -25 | 22 |
| Left MFG 2 | 43 | -34 | 14 |
| Right IPS 2 | -32 | 53 | 49 |
| Right frontal | -44 | -9 | 28 |
| *Right frontoparietal cluster* |  |  |  |
| Right IPS 1 | -44 | 45 | 50 |
| Right IFG | -53 | -15 | 16 |
| Right MFG 1 | -44 | -32 | 30 |
| Right MFG 2 | -43 | -42 | 18 |
| *Sensory cluster (control)* |  |  |  |
| Right cuneus | 19 | 102 | -4 |
| Left cuneus | -16 | 102 | -5 |
| Right lingual gyrus | -1 | 95 | -13 |
| Left lingual gyrus | 10 | 102 | -12 |
| R middle occipital gyrus | -30 | 81 | 14 |

AI = anterior insula; FO = frontal/operculum; IFG = inferior frontal gyrus; IPS = inferior parietal sulcus; MFG = middle frontal gyrus; SMA = supplemental motor area

**Table S2.** Trial Counts by Emotional Expression.

| Emotion/Judgment | Minimum | Maximum | Mean |
| --- | --- | --- | --- |
| Positive Happy | 35 | 60 | 55 |
| Negative Happy | 0 | 24 | 4 |
| Positive Angry | 0 | 17 | 6 |
| Negative Angry | 33 | 59 | 52 |
| Positive Fearful | 1 | 26 | 8 |
| Negative Fearful | 23 | 59 | 50 |
| Positive Surprised | 0 | 20 | 7 |
| Negative Surprised | 32 | 60 | 51 |

Note: There were 60 total trials of each emotional expression.

**Table S3.** Pairwise RT comparisons for the main study.

|  | Emotion | Difference | 95% CI | S.E. | p-value |
| --- | --- | --- | --- | --- | --- |
| *Noise Phase* |  |  |  |  |  |
| **Happy** | **Angry** | **-714.2** | **(-886.1, -542.3)** | **84.2** | **<.001** |
| **Happy** | **Fearful** | **-508.4** | **(-673.0, -343.9)** | **80.6** | **<.001** |
| **Happy** | **Positive Surprised** | **-417.4** | **(-776.1, -58.7)** | **175.6** | **.024** |
| Happy | Negative Surprised | -148.3 | (-356.8, 60.1) | 102.1 | .156 |
| **Angry** | **Fearful** | **205.8** | **(104.1, 307.5)** | **49.8** | **<.001** |
| Angry | Positive Surprised | 296.9 | (-64.0, 657.7) | 176.7 | .103 |
| **Angry** | **Negative Surprised** | **565.9** | **(426.4, 705.4)** | **68.3** | **<.001** |
| Fearful | Positive Surprised | 91.1 | (-249.7, 431.9) | 166.9 | .589 |
| **Fearful** | **Negative Surprised** | **360.1** | **(254.0, 466.2)** | **51.9** | **<.001** |
| Positive Surprised | Negative Surprised | 269.0 | (-69.6, 607.7) | 165.8 | .115 |
| *Reveal Phase* |  |  |  |  |  |
| **Happy** | **Angry** | **-75.7** | **(-93.8, -57.7)** | **8.8** | **<.001** |
| **Happy** | **Fearful** | **-84.2** | **(-111.7, -56.7)** | **13.4** | **<.001** |
| **Happy** | **Positive Surprised** | **-387.4** | **(-450.6, -324.3)** | **30.7** | **<.001** |
| **Happy** | **Negative Surprised** | **-142.5** | **(-187.6, -97.3)** | **22.0** | **<.001** |
| Angry | Fearful | -8.5 | (-26.4, 9.5) | 8.7 | .341 |
| **Angry** | **Positive Surprised** | **-311.7** | **(-368.3, -255.2)** | **27.5** | **<.001** |
| **Angry** | **Negative Surprised** | **-66.8** | **(-106.5, -27.0)** | **19.3** | **.002** |
| **Fearful** | **Positive Surprised** | **-303.3** | **(-358.3, -248.3)** | **26.7** | **<.001** |
| **Fearful** | **Negative Surprised** | **-58.3** | **(-95.1, -21.5)** | **17.9** | **.003** |
| **Positive Surprised** | **Negative Surprised** | **245.0** | **(181.5, 308.4)** | **30.9** | **<.001** |

Note: Bold font indicates a significant difference ( $p < .05$ ).

**Table S4.** Pairwise RT comparisons for the follow-up study.

| Emotion |  | Difference | 95% CI | S.E. | p-value |
| --- | --- | --- | --- | --- | --- |
| *Noise Phase* |  |  |  |  |  |
| **Happy** | **Angry** | **-704.5** | **(-844.0, -564.9)** | **70.1** | **<.001** |
| **Happy** | **Fearful** | **-356.3** | **(-501.3, -211.2)** | **72.9** | **<.001** |
| Happy | Positive Surprised | -180.5 | (-477.8, 116.7) | 149.4 | .230 |
| Happy | Negative Surprised | -122.2 | (-278.7, 34.3) | 78.6 | .124 |
| **Angry** | **Fearful** | **348.2** | **(215.6, 480.9)** | **66.6** | **<.001** |
| **Angry** | **Positive Surprised** | **524.0** | **(217.5, 725.6)** | **154.0** | **.001** |
| **Angry** | **Negative Surprised** | **582.3** | **(438.9, 501.3)** | **72.0** | **<.001** |
| Fearful | Positive Surprised | 175.7 | (-111.6, 463.1) | 144.4 | .227 |
| **Fearful** | **Negative Surprised** | **234.0** | **(105.8, 362.3)** | **64.4** | **<.001** |
| Positive Surprised | Negative Surprised | 58.3 | (-255.6, 372.2) | 157.7 | .713 |
| *Reveal Phase* |  |  |  |  |  |
| **Happy** | **Angry** | **-48.9** | **(-71.0, -26.8)** | **11.1** | **<.001** |
| **Happy** | **Fearful** | **-86.9** | **(-107.1, -66.7)** | **10.1** | **<.001** |
| **Happy** | **Positive Surprised** | **-336.3** | **(-391.0, -281.5)** | **27.5** | **<.001** |
| **Happy** | **Negative Surprised** | **-95.9** | **(-127.6, -64.1)** | **15.9** | **<.001** |
| **Angry** | **Fearful** | **-38.0** | **(-54.7, -21.2)** | **8.4** | **<.001** |
| **Angry** | **Positive Surprised** | **-287.3** | **(-341.8, -232.9)** | **27.3** | **<.001** |
| **Angry** | **Negative Surprised** | **-46.9** | **(-78.8, -15.1)** | **16.0** | **.004** |
| **Fearful** | **Positive Surprised** | **-249.4** | **(-303.5, -195.2)** | **27.2** | **<.001** |
| Fearful | Negative Surprised | -9.0 | (-36.1, 18.2) | 13.6 | .512 |
| **Positive Surprised** | **Negative Surprised** | **240.4** | **(180.4, 300.4)** | **30.1** | **<.001** |

Note: Bold font indicates a significant difference ( $p < .05$ ).

## HRF Emotional Expression Analyses

Linear Mixed Models [Emotion reference = Negative Surprised]

**Table S5.** CO ROI Cluster Mixed Model.

Tests of Fixed Effects:
|  | df | F | p |
| --- | --- | --- | --- |
| **Intercept** | **1, 318.669** | **3.908** | **.049** |
| **Emotion** | **4, 306.265** | **7.918** | **<.001** |
| Time (linear) | 1, 679.043 | 3.350 | .068 |
| **Time (quadratic)** | **1, 696.416** | **49.373** | **<.001** |
| **Emotion \* Time** | **4, 372.264** | **19.573** | **<.001** |
| **Reveal RT** | **1, 925.871** | **65.655** | **<.001** |

Estimates of Fixed Effects:
| Parameter | Estimate | SE | df | t | p | 95% CI |
| --- | --- | --- | --- | --- | --- | --- |
| **Intercept** | **-.112** | **.036** | **322.064** | **-3.130** | **.002** | **-.182, -.042** |
| **Happy** | **.077** | **.015** | **324.562** | **5.300** | **<.001** | **.049, .106** |
| **Angry** | **.062** | **.014** | **342.901** | **4.360** | **<.001** | **.034, .091** |
| **Fearful** | **.043** | **.013** | **311.277** | **3.267** | **.001** | **.017, .069** |
| Positive Surprised | .028 | .029 | 177.188 | .954 | .341 | -.030, .085 |
| **Time (linear)** | **.013** | **.003** | **552.516** | **3.895** | **<.001** | **.007, .020** |
| **Time (quadratic)** | **-.002** | **.000** | **696.416** | **-7.027** | **<.001** | **-.002, -.001** |
| **Happy \* Time** | **-.017** | **.002** | **361.973** | **-7.722** | **<.001** | **-.021, -.013** |
| **Angry \* Time** | **-.014** | **.002** | **390.001** | **-6.163** | **<.001** | **-.018, -.009** |
| **Fearful \* Time** | **-.010** | **.002** | **369.662** | **-4.942** | **<.001** | **-.015, -.006** |
| Positive Surprised * Time | .003 | .004 | 167.975 | .780 | .437 | -.005, .011 |
| **Reveal RT** | **.000** | **.000** | **925.871** | **8.103** | **<.001** | **.000, .000** |

Estimate of Random Effects Covariance Parameters:
| Parameter | Estimate | SE | Wald Z | p | 95% CI |
| --- | --- | --- | --- | --- | --- |
| **Intercept (subject) Variance** | **.009** | **.002** | **3.864** | **<.001** | **.005, .015** |

Note: Bold font indicates a significant effect ( $p < .05$ ).

**Table S6.** L FP ROI Cluster Mixed Model.

Tests of Fixed Effects:
|  | df | F | p |
| --- | --- | --- | --- |
| Intercept | 1, 248.664 | .139 | .710 |
| **Emotion** | **4, 300.360** | **3.634** | **.007** |
| **Time (linear)** | **1, 640.706** | **17.136** | **<.001** |
| **Time (quadratic)** | **1, 713.639** | **77.132** | **<.001** |
| **Emotion \* Time** | **4, 350.836** | **5.095** | **<.001** |
| **Reveal RT** | **1, 904.890** | **16.900** | **<.001** |

Estimates of Fixed Effects:
| Parameter | Estimate | SE | df | t | p | 95% CI |
| --- | --- | --- | --- | --- | --- | --- |
| Intercept | -.009 | .036 | 256.566 | -.257 | .797 | -.079, .061 |
| **Happy** | **.040** | **.015** | **340.395** | **2.689** | **.008** | **.011, .069** |
| **Angry** | **.038** | **.013** | **302.590** | **2.797** | **.005** | **.011, .064** |
| Fearful | .005 | .013 | 283.211 | .413 | .680 | -.020, .031 |
| Positive Surprised | .029 | .026 | 178.416 | 1.092 | .276 | -.023, .080 |
| **Time (linear)** | **.015** | **.003** | **487.322** | **4.757** | **<.001** | **.009, .022** |
| **Time (quadratic)** | **-.002** | **.000** | **713.639** | **-8.782** | **<.001** | **-.003, -.002** |
| **Happy \* Time** | **-.008** | **.002** | **388.469** | **-3.620** | **<.001** | **-.013, -.004** |
| **Angry \* Time** | **-.006** | **.002** | **381.373** | **-2.965** | **.003** | **-.011, -.002** |
| Fearful * Time | -.003 | .002 | 387.589 | -1.403 | .161 | -.007, .001 |
| Positive Surprised * Time | .002 | .004 | 171.487 | .614 | .540 | -.005, .009 |
| **Reveal RT** | **.000** | **.000** | **904.890** | **4.111** | **<.001** | **.000, .000** |

Estimate of Random Effects Covariance Parameters:
| Parameter | Estimate | SE | Wald Z | p | 95% CI |
| --- | --- | --- | --- | --- | --- |
| **Intercept (subject) Variance** | **.011** | **.003** | **3.849** | **<.001** | **.007, .019** |

Note: Bold font indicates a significant effect ( $p < .05$ ).

**Table S7.** R FP ROI Cluster Mixed Model.

Tests of Fixed Effects:
|  | df | F | p |
| --- | --- | --- | --- |
| **Intercept** | **1, 240.547** | **28.876** | **<.001** |
| **Emotion** | **4, 281.994** | **15.618** | **<.001** |
| **Time (linear)** | **1, 630.802** | **431.905** | **<.001** |
| **Time (quadratic)** | **1, 716.230** | **507.716** | **<.001** |
| **Emotion \* Time** | **4, 353.784** | **11.907** | **<.001** |
| **Reveal RT** | **1, 862.805** | **57.919** | **<.001** |

Estimates of Fixed Effects:
| Parameter | Estimate | SE | df | t | p | 95% CI |
| --- | --- | --- | --- | --- | --- | --- |
| **Intercept** | **-.223** | **.038** | **247.748** | **-5.828** | **<.001** | **-.298, -.147** |
| **Happy** | **.105** | **.015** | **324.014** | **6.964** | **<.001** | **.075, .134** |
| **Angry** | **.029** | **.013** | **278.442** | **2.148** | **.033** | **.002, .055** |
| **Fearful** | **.039** | **.013** | **277.223** | **2.876** | **.004** | **.012, .065** |
| **Positive Surprised** | **-.071** | **.026** | **165.515** | **-2.707** | **.008** | **-.123, -.019** |
| **Time (linear)** | **.067** | **.003** | **445.112** | **19.106** | **<.001** | **.060, .073** |
| **Time (quadratic)** | **-.006** | **.000** | **716.230** | **-22.533** | **<.001** | **-.007, -.005** |
| **Happy \* Time** | **-.011** | **.002** | **410.815** | **-4.480** | **<.001** | **-.016, -.006** |
| Angry * Time | -.003 | .002 | 394.083 | -1.419 | .157 | -.008, .001 |
| Fearful * Time | -.004 | .002 | 389.169 | -1.677 | .094 | -.008, .001 |
| **Positive Surprised \* Time** | **.013** | **.004** | **173.781** | **3.429** | **<.001** | **.006, .020** |
| **Reveal RT** | **.000** | **.000** | **862.805** | **7.610** | **<.001** | **.000, .000** |

Estimate of Random Effects Covariance Parameters:
| Parameter | Estimate | SE | Wald Z | p | 95% CI |
| --- | --- | --- | --- | --- | --- |
| **Intercept (subject) Variance** | **.013** | **.003** | **3.879** | **<.001** | **.008, .022** |

Note: Bold font indicates a significant effect ( $p < .05$ ).

**Table S8.** Sensory ROI Cluster Mixed Model.

Tests of Fixed Effects:
|  | df | F | p |
| --- | --- | --- | --- |
| **Intercept** | **1, 92.594** | **62.540** | **<.001** |
| Emotion | 4, 306.107 | .604 | .660 |
| **Time (linear)** | **1, 594.887** | **742.642** | **<.001** |
| **Time (quadratic)** | **1, 723.891** | **944.137** | **<.001** |
| Emotion * Time | 4, 459.008 | .341 | .850 |
| **Reveal RT** | **1, 681.560** | **7.851** | **.005** |

Estimates of Fixed Effects:
| Parameter | Estimate | SE | df | t | p | 95% CI |
| --- | --- | --- | --- | --- | --- | --- |
| **Intercept** | **.885** | **.109** | **95.892** | **8.129** | **<.001** | **.669, 1.101** |
| Happy | -.049 | .036 | 320.077 | -1.379 | .169 | -.120, .021 |
| Angry | -.041 | .038 | 298.696 | -1.081 | .281 | -.115, .033 |
| Fearful | -.043 | .034 | 275.697 | -1.258 | .210 | -.111, .024 |
| Positive Surprised | -.030 | .063 | 191.277 | -.473 | .637 | -.154, .095 |
| **Time (linear)** | **.228** | **.009** | **382.954** | **25.031** | **<.001** | **.210, .246** |
| **Time (quadratic)** | **-.024** | **.001** | **723.891** | **-30.727** | **<.001** | **-.025, -.022** |
| Happy * Time | .005 | .006 | 471.234 | .761 | .447 | -.008, .017 |
| Angry * Time | .003 | .007 | 481.062 | .433 | .665 | -.011, .016 |
| Fearful * Time | .006 | .007 | 467.424 | .910 | .363 | -.007, .019 |
| Positive Surprised * Time | .010 | .010 | 233.667 | .930 | .353 | -.011, .030 |
| **Reveal RT** | **.000** | **.000** | **681.560** | **-2.802** | **.005** | **.000, .000** |

Estimate of Random Effects Covariance Parameters:
| Parameter | Estimate | SE | Wald Z | p | 95% CI |
| --- | --- | --- | --- | --- | --- |
| **Intercept (subject) Variance** | **.209** | **.053** | **3.920** | **<.001** | **.127, .344** |

Note: Bold font indicates a significant effect ( $p < .05$ ).

**Table S9.**
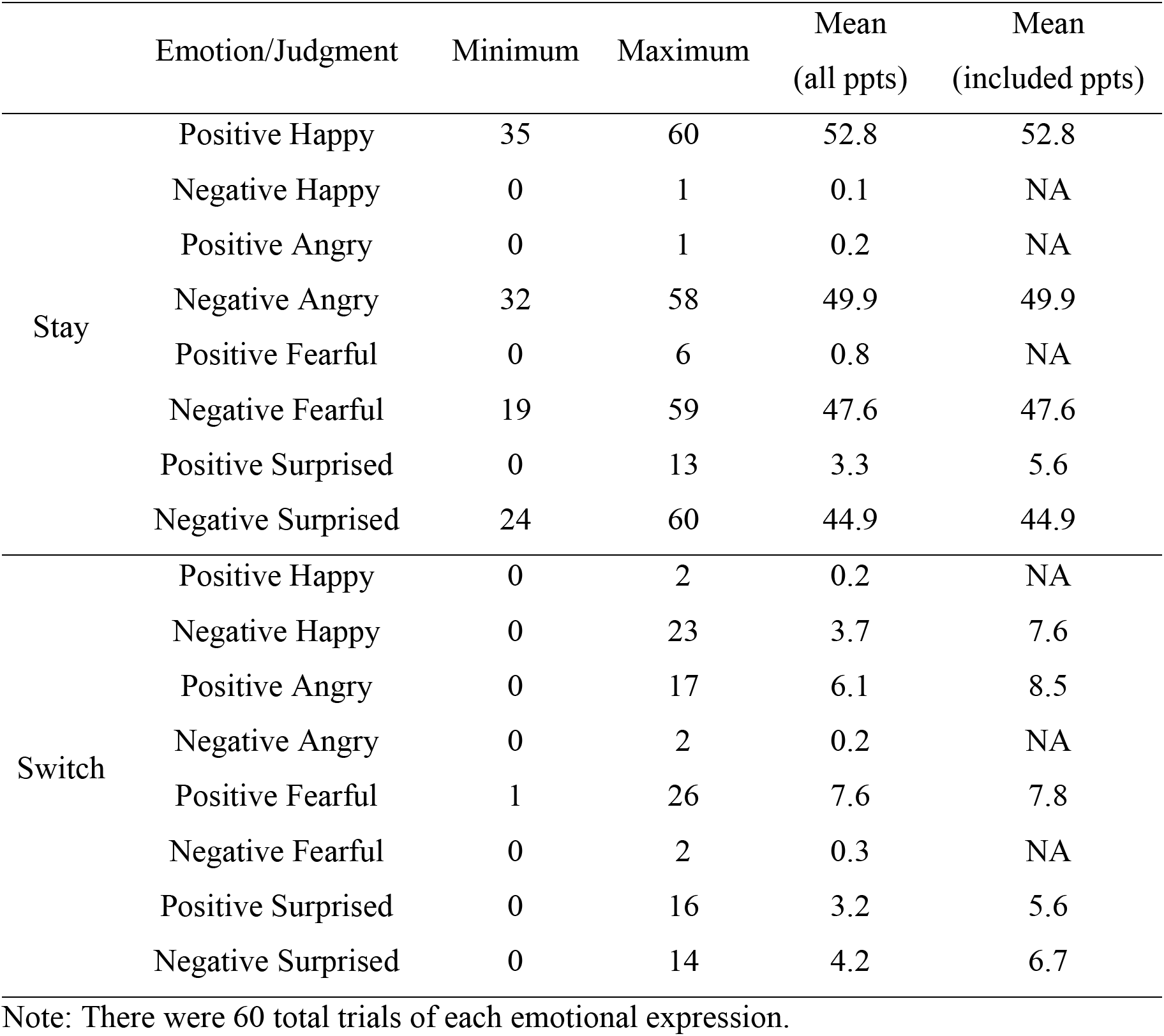
Trial Counts by Stay/Switch.

|  | Emotion/Judgment | Minimum | Maximum | Mean<br>(all ppts) | Mean<br>(included ppts) |
| --- | --- | --- | --- | --- | --- |
| Stay | Positive Happy | 35 | 60 | 52.8 | 52.8 |
|  | Negative Happy | 0 | 1 | 0.1 | NA |
|  | Positive Angry | 0 | 1 | 0.2 | NA |
|  | Negative Angry | 32 | 58 | 49.9 | 49.9 |
|  | Positive Fearful | 0 | 6 | 0.8 | NA |
|  | Negative Fearful | 19 | 59 | 47.6 | 47.6 |
|  | Positive Surprised | 0 | 13 | 3.3 | 5.6 |
|  | Negative Surprised | 24 | 60 | 44.9 | 44.9 |
| Switch | Positive Happy | 0 | 2 | 0.2 | NA |
|  | Negative Happy | 0 | 23 | 3.7 | 7.6 |
|  | Positive Angry | 0 | 17 | 6.1 | 8.5 |
|  | Negative Angry | 0 | 2 | 0.2 | NA |
|  | Positive Fearful | 1 | 26 | 7.6 | 7.8 |
|  | Negative Fearful | 0 | 2 | 0.3 | NA |
|  | Positive Surprised | 0 | 16 | 3.2 | 5.6 |
|  | Negative Surprised | 0 | 14 | 4.2 | 6.7 |

Note: There were 60 total trials of each emotional expression.

**Table S10.** CO ROI Cluster Switch Trials Mixed Model.

Tests of Fixed Effects:
|  | df | F | p |
| --- | --- | --- | --- |
| **Intercept** | **1, 76.644** | **8.388** | **.005** |
| **Emotion** | **4, 199.821** | **26.090** | **<.001** |
| **Time (linear)** | **1, 476.728** | **181.641** | **<.001** |
| **Time (quadratic)** | **1, 510.659** | **240.315** | **<.001** |
| **Emotion \* Time** | **4, 204.771** | **45.333** | **<.001** |

Estimates of Fixed Effects:
| Parameter | Estimate | SE | df | t | p | 95% CI |
| --- | --- | --- | --- | --- | --- | --- |
| **Intercept** | **-.079** | **.037** | **113.871** | **-2.169** | **.032** | **-.152, -.007** |
| **Happy** | **.258** | **.037** | **177.630** | **6.974** | **<.001** | **.185, .331** |
| **Angry** | **.277** | **.030** | **176.152** | **9.388** | **<.001** | **.219, .335** |
| **Fearful** | **.204** | **.031** | **204.402** | **6.631** | **<.001** | **.143, .265** |
| **Positive Surprised** | **.088** | **.042** | **162.642** | **2.096** | **.038** | **.005, .170** |
| **Time (linear)** | **.129** | **.007** | **265.246** | **18.084** | **<.001** | **.115, .143** |
| **Time (quadratic)** | **-.008** | **.001** | **510.659** | **-15.502** | **<.001** | **-.009, -.007** |
| **Happy \* Time** | **-.062** | **.006** | **188.119** | **-10.506** | **<.001** | **-.073, -.050** |
| **Angry \* Time** | **-.059** | **.005** | **170.156** | **-12.773** | **<.001** | **-.068, -.050** |
| **Fearful \* Time** | **-.050** | **.005** | **193.080** | **-10.321** | **<.001** | **-.059, -.040** |
| **Positive Surprised \* Time** | **-.037** | **.006** | **174.004** | **-5.767** | **<.001** | **-.049, -.024** |

Estimate of Random Effects Covariance Parameters:
| Parameter | Estimate | SE | Wald Z | p | 95% CI |
| --- | --- | --- | --- | --- | --- |
| **Intercept (subject) Variance** | **.017** | **.004** | **3.759** | **<.001** | **.010, .028** |

Note: Bold font indicates a significant effect ( $p < .05$ ).

**Table S11.** CO ROI Cluster Stay Trials Mixed Model.

Tests of Fixed Effects:
|  | df | F | p |
| --- | --- | --- | --- |
| **Intercept** | **1, 57.877** | **74.890** | **<.001** |
| **Emotion** | **4, 220.562** | **6.794** | **<.001** |
| Time (linear) | 1, 687.945 | 1.860 | .173 |
| **Time (quadratic)** | **1, 684.896** | **42.716** | **<.001** |
| **Emotion \* Time** | **4, 281.603** | **16.205** | **<.001** |

Estimates of Fixed Effects:
| Parameter | Estimate | SE | df | t | p | 95% CI |
| --- | --- | --- | --- | --- | --- | --- |
| **Intercept** | **.108** | **.019** | **64.923** | **5.671** | **<.001** | **.070, .147** |
| **Happy** | **.037** | **.014** | **273.689** | **2.680** | **.008** | **.010, .064** |
| **Angry** | **.051** | **.014** | **333.131** | **3.681** | **<.001** | **.024, .079** |
| **Fearful** | **.031** | **.013** | **301.785** | **2.452** | **.015** | **.006, .056** |
| **Positive Surprised** | **.137** | **.032** | **90.230** | **4.306** | **<.001** | **.074, .201** |
| **Time (linear)** | **.009** | **.003** | **528.267** | **2.631** | **.009** | **.002, .015** |
| **Time (quadratic)** | **-.002** | **.000** | **684.896** | **-6.536** | **<.001** | **-.002, -.001** |
| **Happy \* Time** | **-.014** | **.002** | **349.255** | **-6.249** | **<.001** | **-.018, -.009** |
| **Angry \* Time** | **-.012** | **.002** | **384.360** | **-5.330** | **<.001** | **-.016, -.007** |
| **Fearful \* Time** | **-.009** | **.002** | **352.404** | **-4.172** | **<.001** | **-.013, -.005** |
| **Positive Surprised \* Time** | **.012** | **.005** | **114.409** | **2.520** | **.013** | **.003, .022** |

Estimate of Random Effects Covariance Parameters:
| Parameter | Estimate | SE | Wald Z | p | 95% CI |
| --- | --- | --- | --- | --- | --- |
| **Intercept (subject) Variance** | **.008** | **.002** | **3.858** | **<.001** | **.005, .013** |

Note: Bold font indicates a significant effect ( $p < .05$ ).

**Figure S2.**
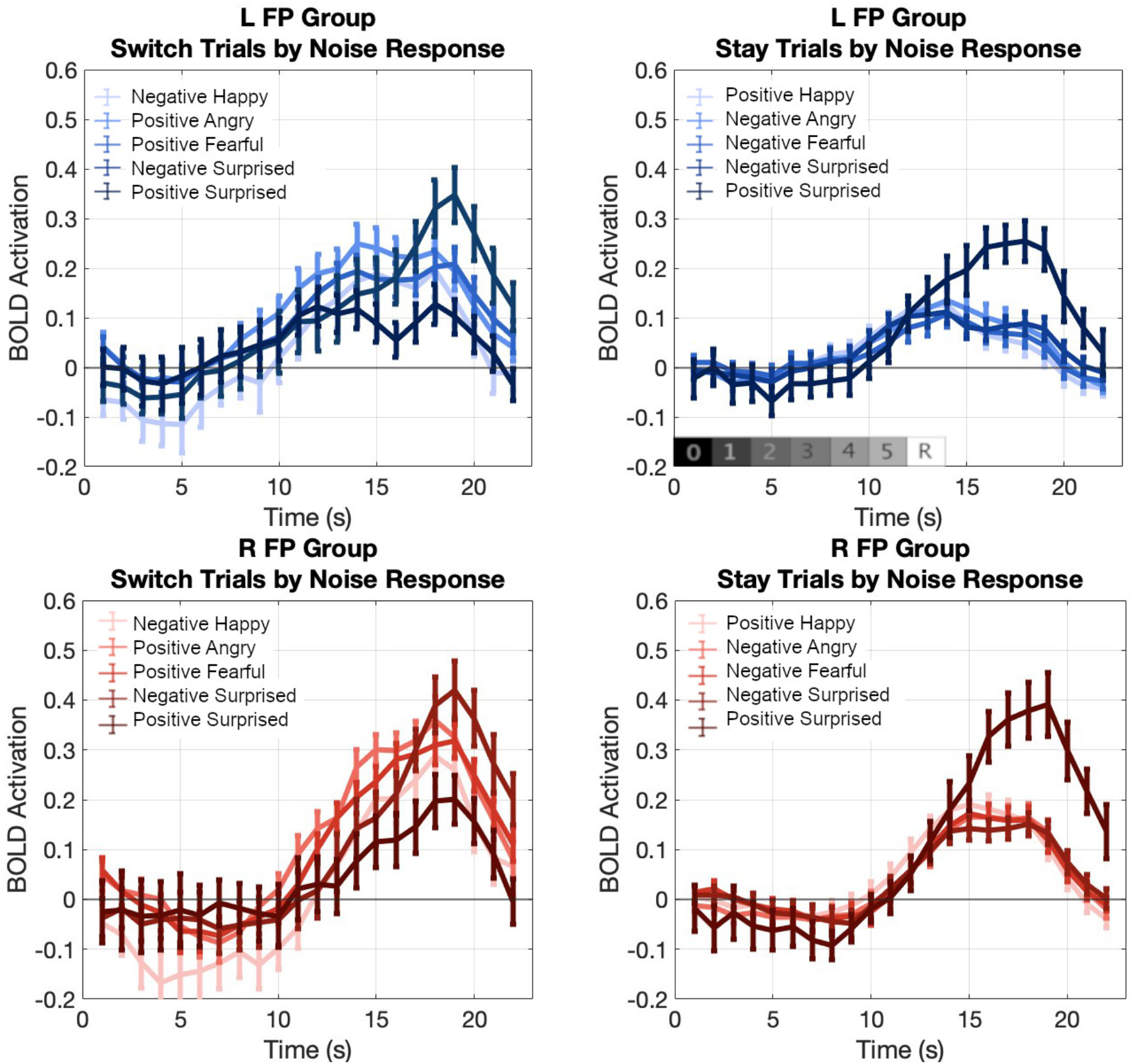
Estimated HRF curves in the L FP and R FP clusters for each emotion based on whether a participant switched their behavioral response from the noise to the reveal phase or stayed with the same response. Labels refer to the initial response made during the noise phase.

**Table S12.**
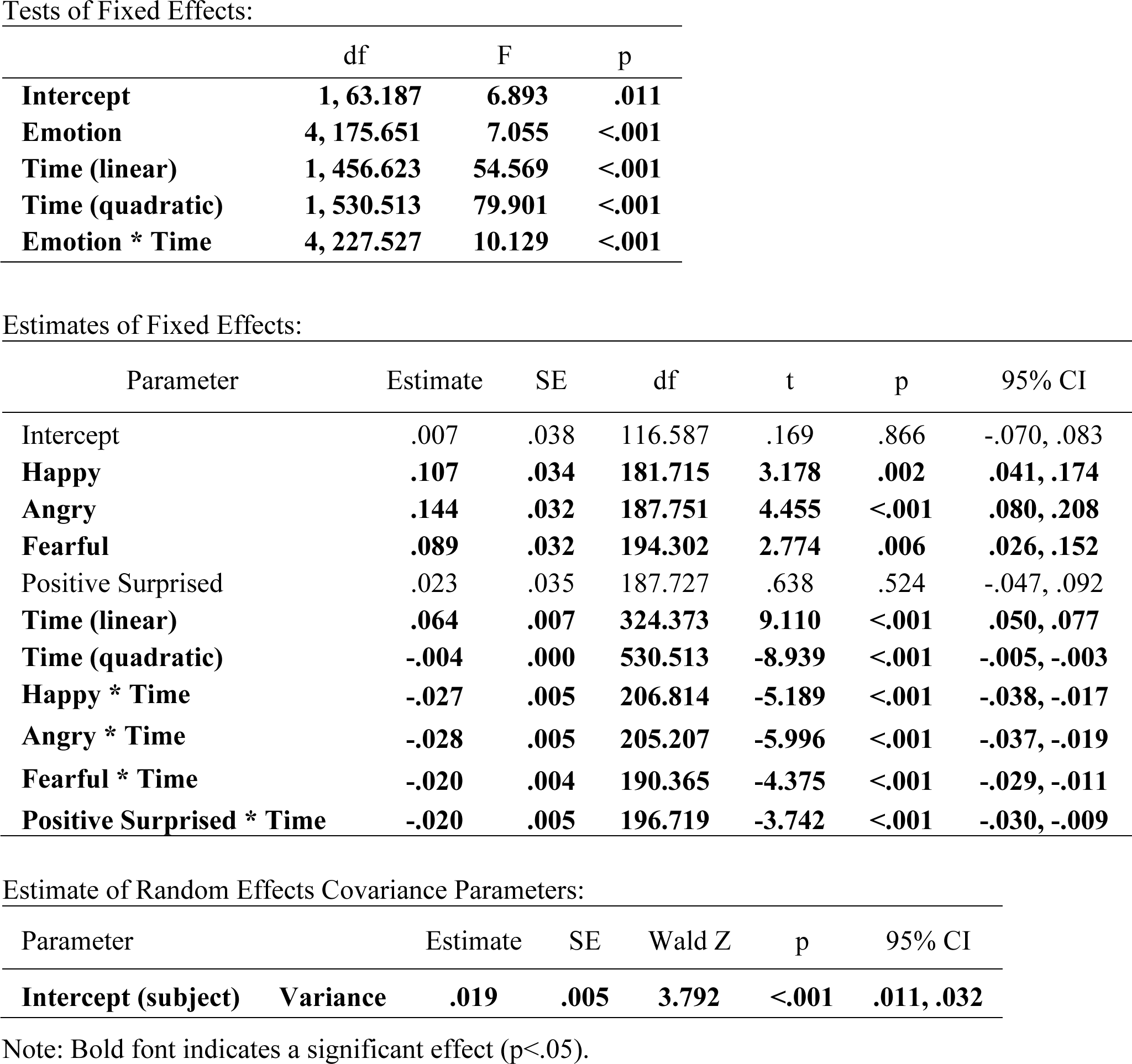
L FP ROI Cluster Switch Trials Mixed Model.

**Table S13.** L FP ROI Cluster Stay Trials Mixed Model.

Tests of Fixed Effects:
|  | df | F | p |
| --- | --- | --- | --- |
| **Intercept** | **1, 43.474** | **26.381** | **<.001** |
| **Emotion** | **4, 249.565** | **5.998** | **<.001** |
| **Time (linear)** | **1, 645.773** | **10.403** | **.001** |
| **Time (quadratic)** | **1, 708.775** | **60.634** | **<.001** |
| **Emotion \* Time** | **4, 327.337** | **3.982** | **.004** |

Estimates of Fixed Effects:
| Parameter | Estimate | SE | df | t | p | 95% CI |
| --- | --- | --- | --- | --- | --- | --- |
| **Intercept** | **.082** | **.023** | **51.096** | **3.514** | **<.001** | **.035, .129** |
| Happy | .025 | .014 | 297.207 | 1.774 | .077 | -.003, .054 |
| **Angry** | **.034** | **.014** | **296.794** | **2.523** | **.012** | **.008, .061** |
| Fearful | -.001 | .013 | 287.483 | -.113 | .910 | -.027, .024 |
| **Positive Surprised** | **.105** | **.027** | **102.964** | **3.926** | **<.001** | **.052, .158** |
| **Time (linear)** | **.012** | **.003** | **479.340** | **3.613** | **<.001** | **.005, .018** |
| **Time (quadratic)** | **-.002** | **.000** | **708.775** | **-7.787** | **<.001** | **-.002, -.001** |
| **Happy \* Time** | **-.007** | **.002** | **380.136** | **-2.986** | **.003** | **-.011, -.002** |
| **Angry \* Time** | **-.005** | **.002** | **380.071** | **-2.268** | **.024** | **-.009, -.001** |
| Fearful * Time | -.002 | .002 | 396.031 | -.810 | .418 | -.006, .003 |
| Positive Surprised * Time | .004 | .004 | 125.697 | 1.036 | .302 | -.004, .012 |

Estimate of Random Effects Covariance Parameters:
| Parameter | Estimate | SE | Wald Z | p | 95% CI |
| --- | --- | --- | --- | --- | --- |
| **Intercept (subject) Variance** | **.013** | **.003** | **3.880** | **<.001** | **.008, .022** |

Note: Bold font indicates a significant effect ( $p < .05$ ).

**Table S14.** R FP ROI Cluster Switch Trials Mixed Model.

Tests of Fixed Effects:
|  | df | F | p |
| --- | --- | --- | --- |
| **Intercept** | **1, 75.156** | **4.379** | **.040** |
| **Emotion** | **4, 189.618** | **12.624** | **<.001** |
| **Time (linear)** | **1, 429.312** | **250.535** | **<.001** |
| **Time (quadratic)** | **1, 511.999** | **215.660** | **<.001** |
| **Emotion \* Time** | **4, 240.188** | **10.233** | **<.001** |

Estimates of Fixed Effects:
| Parameter | Estimate | SE | df | t | p | 95% CI |
| --- | --- | --- | --- | --- | --- | --- |
| **Intercept** | **-.162** | **.037** | **122.624** | **-4.336** | **<.001** | **-.235, -.088** |
| **Happy** | **.109** | **.038** | **171.729** | **2.866** | **.005** | **.034, .185** |
| **Angry** | **.199** | **.030** | **186.158** | **6.514** | **<.001** | **.138, .259** |
| **Fearful** | **.148** | **.031** | **201.777** | **4.760** | **<.001** | **.087, .210** |
| Positive Surprised | .046 | .041 | 146.697 | 1.099 | .274 | -.036, .127 |
| **Time (linear)** | **.125** | **.008** | **262.559** | **16.492** | **<.001** | **.110, .140** |
| **Time (quadratic)** | **-.008** | **.001** | **511.999** | **-14.685** | **<.001** | **-.009, -.007** |
| **Happy \* Time** | **-.024** | **.006** | **196.352** | **-3.877** | **<.001** | **-.036, -.012** |
| **Angry \* Time** | **-.031** | **.005** | **219.684** | **-6.239** | **<.001** | **-.041, -.021** |
| **Fearful \* Time** | **-.025** | **.005** | **220.415** | **-5.080** | **<.001** | **-.035, -.015** |
| **Positive Surprised \* Time** | **-.022** | **.006** | **169.107** | **-3.598** | **<.001** | **-.035, -.010** |

Estimate of Random Effects Covariance Parameters:
| Parameter | Estimate | SE | Wald Z | p | 95% CI |
| --- | --- | --- | --- | --- | --- |
| **Intercept (subject) Variance** | **.016** | **.004** | **3.759** | **<.001** | **.010, .028** |

Note: Bold font indicates a significant effect ( $p < .05$ ).

**Table S15.** R FP ROI Cluster Stay Trials Mixed Model.

Tests of Fixed Effects:
|  | df | F | p |
| --- | --- | --- | --- |
| Intercept | 1, 41.488 | .245 | .623 |
| **Emotion** | **4, 206.444** | **6.282** | **<.001** |
| **Time (linear)** | **1, 633.952** | **421.496** | **<.001** |
| **Time (quadratic)** | **1, 692.027** | **499.357** | **<.001** |
| **Emotion \* Time** | **4, 318.821** | **15.443** | **<.001** |

Estimates of Fixed Effects:
| Parameter | Estimate | SE | df | t | p | 95% CI |
| --- | --- | --- | --- | --- | --- | --- |
| Intercept | -.010 | .024 | 49.353 | -.428 | .671 | -.058, .038 |
| **Happy** | **.068** | **.014** | **269.800** | **4.703** | **<.001** | **.040, .097** |
| Angry | .012 | .013 | 280.765 | .881 | .379 | -.015, .038 |
| Fearful | .020 | .013 | 277.779 | 1.474 | .142 | -.007, .046 |
| Positive Surprised | .009 | .026 | 72.178 | .331 | .741 | -.043, .060 |
| **Time (linear)** | **.064** | **.003** | **448.517** | **18.476** | **<.001** | **.057, .071** |
| **Time (quadratic)** | **-.006** | **.000** | **692.027** | **-22.346** | **<.001** | **-.007, -.005** |
| **Happy \* Time** | **-.009** | **.003** | **402.263** | **-3.630** | **<.001** | **-.014, -.004** |
| Angry * Time | -.001 | .002 | 389.610 | -.240 | .811 | -.005, .004 |
| Fearful * Time | -.002 | .002 | 385.648 | -.798 | .425 | -.006, .003 |
| **Positive Surprised \* Time** | **.023** | **.004** | **118.870** | **5.433** | **<.001** | **.015, .031** |

Estimate of Random Effects Covariance Parameters:
| Parameter | Estimate | SE | Wald Z | p | 95% CI |
| --- | --- | --- | --- | --- | --- |
| **Intercept (subject) Variance** | **.014** | **.004** | **3.879** | **<.001** | **.009, .024** |

Note: Bold font indicates a significant effect ( $p < .05$ ).

**Figure S3.**
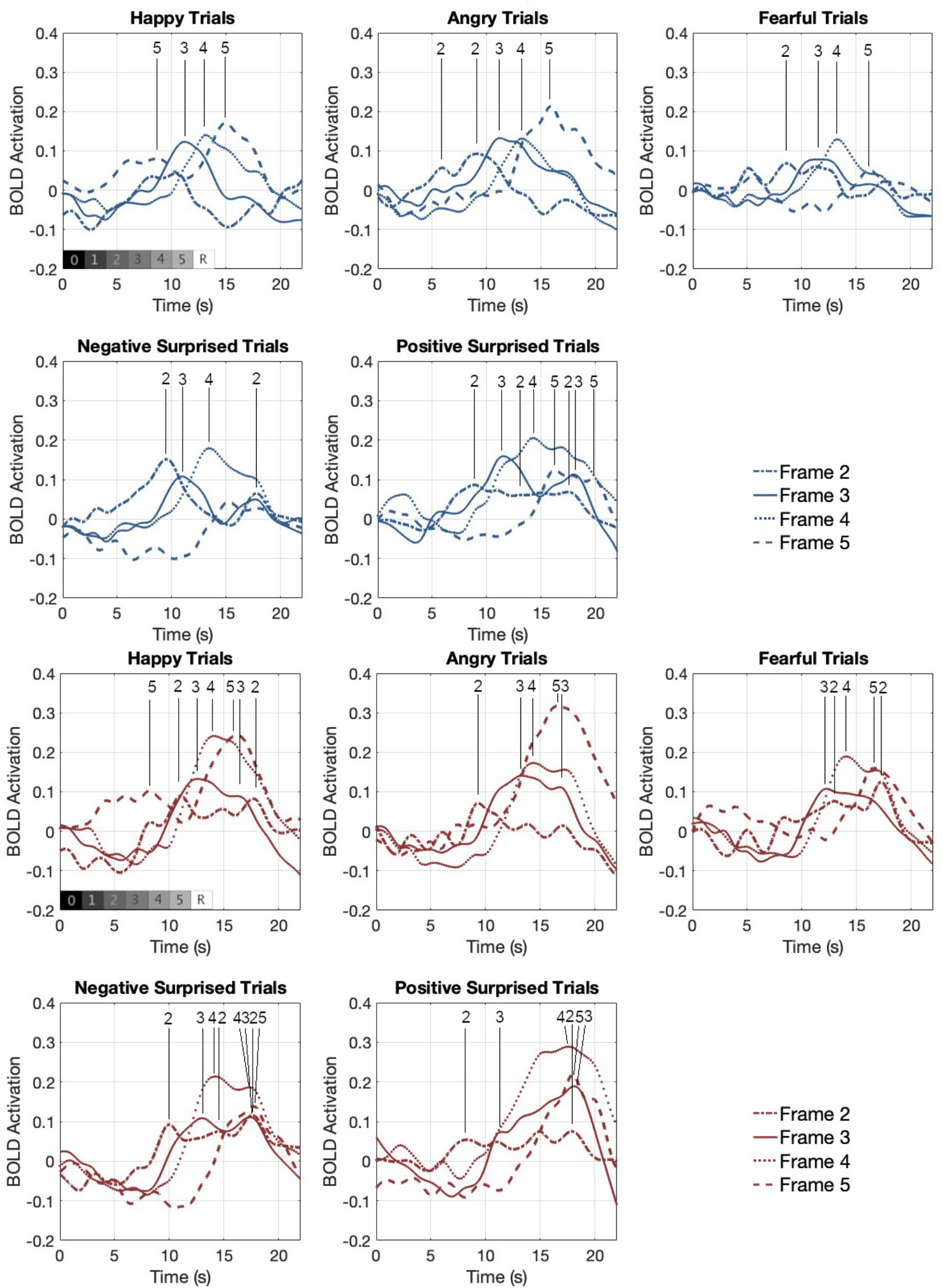
Estimated HRF curves for each emotional expression by response frame for the L FP (blue) and R FP (red) ROI clusters. Averaged responses across participants according to the noise frame (2-5) during which a behavioral response was made for correct clear valence trials (happy, angry, fearful) and ambiguous valence (surprised) trials for negative and positive responses. Numbers at the top of each plot indicate the identified peak(s) for each frame.

**Table S16.** Peak times (seconds) for each ROI Cluster by Emotional Expression and Frame.

| Cluster | Emotion | Frame 2 | Frame 3 | Frame 4 | Frame 5 |
| --- | --- | --- | --- | --- | --- |
| CO | Happy | 9.7 | 11.3 | 13.0 | 14.9 |
|  | Angry | 9.5 | 11.4 | 13.0 | 15.8 |
|  | Fearful | 8.9 | 11.3 | 13.2 | 16.0 |
|  | Negative Surprised | 9.6, 17.7 | 11.1, 17.6 | 13.1, 17.4 | 15.9 |
|  | Positive Surprised | 10.0, 18.1 | 12.4, 17.2 | 16.0 | 18.0 |
| L FP | Happy | --- | 11.2 | 13.1 | 8.6, 15.0 |
|  | Angry | 5.9, 9.1 | 11.2 | 13.2 | 15.9 |
|  | Fearful | 8.6 | 11.3 | 13.3 | 16.2 |
|  | Negative Surprised | 9.5, 17.8 | 11.0 | 13.4 | --- |
|  | Positive Surprised | 8.9, 13.1, 17.4 | 11.5, 18.0 | 14.2 | 16.2, 19.8 |
| R FP | Happy | 11.0, 17.7 | 12.6, 16.2 | 14.0 | 8.1, 16.2 |
|  | Angry | 9.3 | 13.3, 16.8 | 14.4 | 16.6 |
|  | Fearful | 13.0, 17.2 | 12.2 | 14.1 | 16.6 |
|  | Negative Surprised | 10.0, 14.6, 17.6 | 13.0, 17.4 | 14.2, 17.3 | 17.7 |
|  | Positive Surprised | 8.3, 17.9 | 11.5, 18.1 | 17.6 | 18.0 |

